# VAP-A intrinsically disordered regions enable versatile tethering at membrane contact sites

**DOI:** 10.1101/2022.05.13.491839

**Authors:** Mélody Subra, Manuela Dezi, Joëlle Bigay, Sandra Lacas-Gervais, Aurélie Di Cicco, Ana Rita Dias Araújo, Sophie Abélanet, Lucile Fleuriot, Delphine Debayle, Romain Gautier, Amanda Patel, Fanny Roussi, Bruno Antonny, Daniel Lévy, Bruno Mesmin

## Abstract

Membrane contact sites (MCSs) between organelles are heterogeneous in shape, composition and dynamics. Despite this diversity, VAP proteins act as receptors for multiple FFAT motif-containing proteins and drive the formation of most MCSs involving the endoplasmic reticulum (ER). Although the VAP‒FFAT interaction is well characterized, no model explains how VAP adapts to its partners in various MCSs. We report here that VAP-A localization to different MCSs depends on its intrinsically disordered regions (IDRs). We show that VAP-A interaction with PTPIP51 and VPS13A at ER‒mitochondria MCS conditions mitochondria fusion by promoting lipid transfer and cardiolipin buildup. VAP-A also enables lipid exchange at ER‒Golgi MCS by interacting with OSBP and CERT. However, removing IDRs from VAP-A restricts its distribution and function to ER‒ mitochondria MCS, at the expense of ER‒Golgi MCS. Our data suggest that IDRs of VAP-A do not modulate its preference towards specific partners, but adjust its geometry to the constraints linked to different MCS organization and lifetime. Thus, VAP-A conformational flexibility mediated by its IDRs ensures membrane tethering plasticity and efficiency.

## INTRODUCTION

A prominent mechanism of communication between subcellular compartments is to bring them in contact. These membrane contact sites (MCSs) have different functions, such as lipid transfer, calcium signaling and organelle dynamics (Wu et al., 2018). MCS dysfunction is involved in a number of human diseases, including cancer, obesity and neurological disorders (Prinz et al., 2020). In addition, many viruses exploit MCS components for their assembly in cells (Wong et al., 2021). MCSs are ubiquitous, but heterogeneous in shape, composition and dynamics, with intermembrane distances ranging from 15 to 45 nm and lifetime of less than a minute to a few hours. For example, evidence suggests that contacts between the ER and the *trans*-Golgi network (TGN) (abbreviated here as ER‒Golgi MCS) are short-lived, because they involve the rapid turnover of several tethering components in the context of continuous TGN membrane remodeling (David et al., 2021; Mesmin et al., 2019). We have previously demonstrated that Oxysterol-binding protein (OSBP) directs cholesterol transport from the ER to the TGN through coupled counter-exchange and hydrolysis of the phosphoinositide phosphatidylinositol-4-phosphate (PI(4)P) at MCS between these two organelles (Mesmin et al., 2013, 2017). Since PI(4)P also serves as an anchor at the TGN surface for several lipid-transfer proteins (LTPs), the lipid exchange activity of OSBP dynamically modulates membrane tethering. Furthermore, recent findings showed that OSBP contains a long N-terminal intrinsically disordered region (IDR), which prevents protein crowding at ER‒Golgi MCS and facilitates lateral protein movements (Jamecna et al., 2019), thereby suggesting that OSBP is also well equipped to meet the transient nature of these contacts.

In contrast to ER‒Golgi MCS, live-cell imaging studies indicate that contacts between the ER and mitochondria (hereafter ER‒Mito MCS) are stable and persist over time despite the dynamics experienced by both these organelles (Friedman et al., 2010; Guo et al., 2018; Lewis et al., 2016; Rowland and Voeltz, 2012). ER‒Mito MCSs are notably well described for their involvement in mitochondria fission and fusion events, by enabling the accumulation of dedicated protein machineries at precise sites between the facing membranes (Abrisch et al., 2020), and by stabilizing mitochondria all along these processes (Guo et al., 2018). Since ER‒Mito MCSs are also hubs for lipid trafficking (Petrungaro and Kornmann, 2019; Vance, 2014), a related question is whether they help shape a membrane environment conductive to fission or fusion reactions.

Despite the heterogeneous structural stability of MCSs, the ubiquitously expressed VAP proteins are common major players for generating tethers between the ER and other compartments (Alli-Balogun and Levine, 2019; Eisenberg-Bord et al., 2016; Wu et al., 2018). VAPs form a family of evolutionarily conserved proteins that are involved in several functions at MCSs, including lipid transport and homeostasis, and organelle trafficking (Dong et al., 2016; Hua et al., 2017; Raiborg et al., 2015; Wijdeven et al., 2016). The abundant ER-anchored type-II membrane protein VAP-A, and its close homolog VAP-B, which point mutations are associated with motor neuron diseases (Dudás et al., 2021), contain an N-terminal major sperm protein (MSP) domain that binds to numerous proteins containing a FFAT (two phenylalanines in an acidic tract) or FFAT-like motif (Loewen et al., 2003; Mattia et al., 2020), a central region with a predicted coiled-coiled domain responsible for dimerization, and a C-terminal transmembrane domain. VAP-A/B are used extensively as anchors by LTPs to create functional complexes that can bridge membranes and exchange lipids between them, such as the VAP‒OSBP complex at ER‒Golgi MCS (Mesmin et al., 2013; Wyles et al., 2002), or the VAP‒protein tyrosine phosphatase-interacting protein 51 (PTPIP51) complex at ER‒Mito MCS (Stoica et al., 2014; Yeo et al., 2021). In addition, VAP-A/B may also bind to multiple interactors within a single MCS (Peretti et al., 2008).

The structural basis of the MSP‒FFAT interaction is well characterized by X-ray and NMR (Furuita et al., 2010; Kaiser et al., 2005), however it is unclear how VAP adapts to a variety of partners having a wide range of structures and functions, and involved in MCSs of contrasting dynamics. We have shown recently that VAP-A is a highly flexible protein that can tune its length to allow the formation of MCSs of adjustable thickness, from 10 to 30 nm, *in vitro* (Mora et al., 2021). Accordingly, it is possible that VAP-A accommodates different partners and/or supports membrane tethering in various MCS contexts through specific molecular properties that extend beyond the MSP‒FFAT interaction model.

Here, we investigate the impact of VAP-A architectural features on the function, dynamics and organization of MCSs. We show that downstream of the MSP domain, VAP-A contains two stretchable IDRs that link its structured domains together. These linkers are responsible for VAP-A flexibility and ensure its engagement and function at different MCSs: VAP-A supports OSBP- and CERT-mediated lipid exchange at ER‒Golgi MCSs, whereas it plays an important and previously uncharacterized role at ER‒Mito MCS by promoting cardiolipin buildup in mitochondria and mitochondrial fusion. By replacing endogenous VAP-A with a construct lacking flexible IDRs, we show that it massively redistributes to ER‒Mito MCSs, thereby affecting lipid flux at ER‒Golgi MCS. We demonstrate that this shift is due to the incompatibility between a rigid tether and the high turnover experienced by ER‒Golgi MCSs. This study provides new evidence to define VAP as a general and versatile ER receptor and describes IDRs as key features enabling this function at MCS.

## RESULTS

### VAP-A linkers are intrinsically disordered regions

We previously demonstrated that full-length VAP-A reconstituted into liposomes extends its MSP to varying heights from the membrane depending on protein density (Mora et al., 2021). Using in vitro reconstituted MCSs, we also showed that VAP-A conformational change is coupled to the size of its interactor, i.e., VAP-A is tilted when in complex with OSBP, but exhibits a straighter conformation with a shorter OSBP construct, N-PH-FFAT, lacking the bulky OSBP-related domain (ORD). These data suggest that VAP-A is well equipped to adapt vertically to a large spectrum of interactors of different structures (Mora et al., 2021). In this study, we first sought to define the molecular features that provide VAP-A with flexibility. We noticed that downstream to its MSP, VAP-A contains two linker regions (hereafter termed L1 and L2) of 32 and 24 residues, flanking its coiled-coil domain (Fig. 1a). These sequences show lower amino acid conservation than the MSP, coiled-coil and TM domains, but are consistently present across species (Fig. S1a). Different algorithms predict VAP-A linkers to be intrinsically disordered regions (IDRs), which is also suggested by the AlphaFold structure prediction (Fig. S1a,b). Additionally, by analyzing the behavior of these linkers using molecular dynamics (MD) simulations, we found that they were extremely mobile and showed large bending capacity (Fig. S1a,b).

**Fig. 1.**
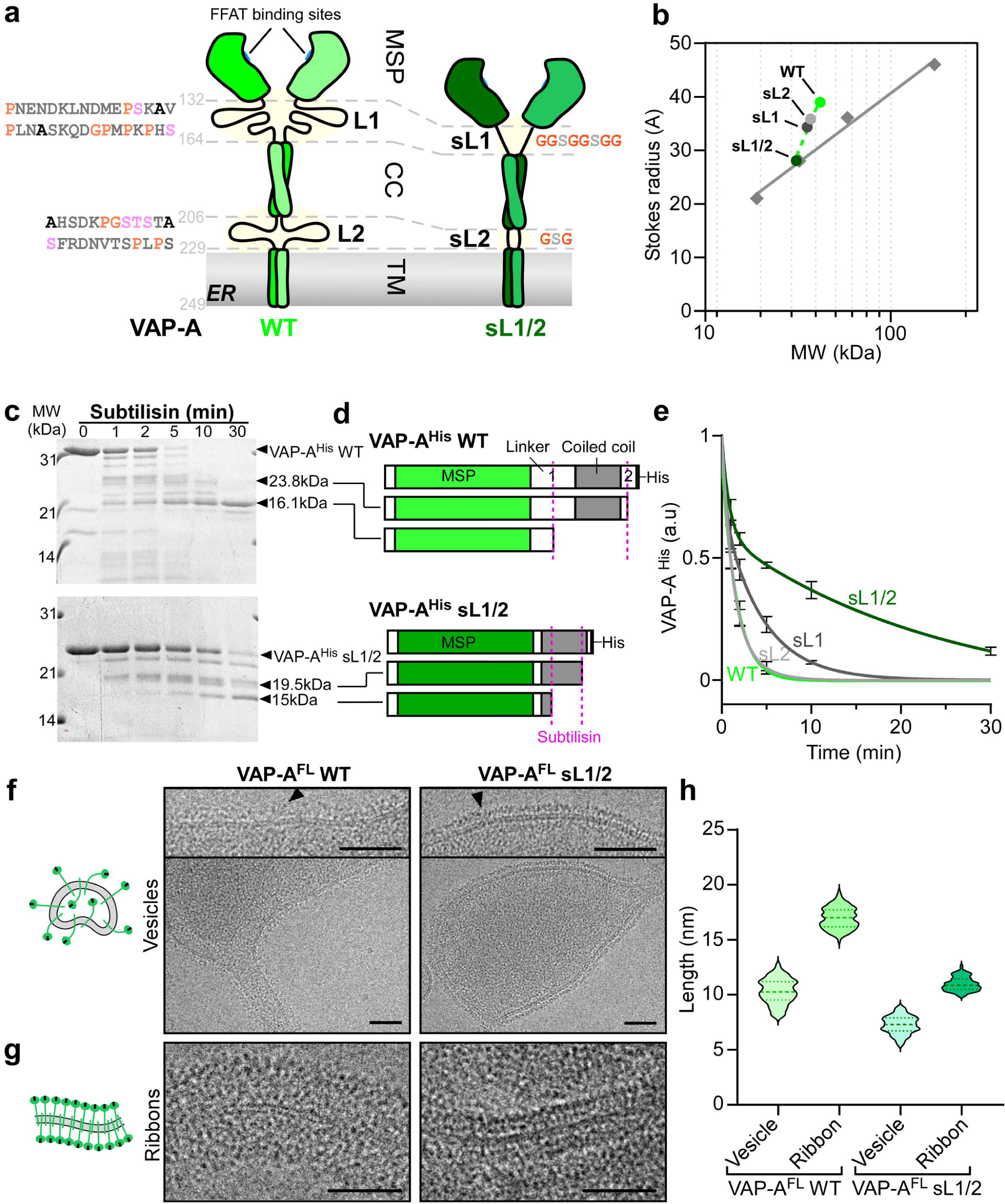
VAP-A linkers are intrinsically disordered regions responsible for its flexibility. **a**, Domain organization of human VAP-A featuring linker regions (L1, L2) with detailed amino acid sequences. The mutant construct with short linker 1 and 2 (VAP-A sL1/2) used in this study is depicted on the right. **b**, Gel filtration analysis of VAP-A^His^ constructs with WT, sL1, sL2 or sL1/sL2 linkers. Stokes radius versus molecular mass (MW) of VAP-A constructs (circles) and folded standards (squares). The dependence of Stokes radius values on the relative molecular masses of VAP-A constructs deviates from that of globular proteins, indicating that VAP-A linkers are intrinsically unfolded. Note that the interception of the two lines coincides with the sL1/2 construct. **c**, Subtilisin digestion of VAP-A^His^ WTor sL1/2 analyzed by SDS-PAGE and Sypro-orange staining. Arrowheads indicate major VAP-A fragments identified by mass spectrometry, also depicted in **d**. VAP-A sL1/2 shows a stronger resistance to Subtilisin cleavage. **e**, Fraction of VAP-A^His^ constructs remaining upon Subtilisin addition over time. The graph represents means of 3 independent experiments ± SEM. **f, g**, Representative cryo-EM images of reconstituted VAP-A^FL^ WT or sL1/2 into proteoliposomes at high protein density (lipid/protein ratio of 70 mol/mol) leading to the formation of spherical or slightly deformed vesicles (f), or ribbon-like membrane fragments with proteins distributed on both sides of the bilayer (g). Zoomed views of electron densities corresponding to VAP-A WT or sL1/2 (arrowheads) at vesicle surface are shown in insets. Bars: 50 nm. **h**, Violin plots showing the length distribution of the extramembrane domains of VAP-A. The horizontal dashed lines show the average values and standard deviations. 19 vesicles (*n*=122 measurements) and 15 ribbons (*n*= 119 measurements) with VAP-A WT were analyzed; 30 vesicles (*n*= 131 measurements) and 14 ribbons (*n*= 113 measurements) with VAP-A sL1/2 were analyzed.

To gain insight into the conformational properties provided by VAP-A linkers, we built several VAP-A constructs in which we replaced the original linkers by much shorter ones, comprising only GGSGGSGG (8 aa., sL1) and GSG (3 aa., sL2) (Fig. 1a, see Methods). Considering that IDRs have larger hydrodynamic dimensions than folded domains of the same amino acid length, we first examined the influence of the two linkers on protein size by gel filtration chromatography (Fig. 1b). To simplify the analysis, we used VAP-A constructs with a 6×His-tag replacing the C-terminal TM domain (VAP^His^ WT, sL1, sL2 and sL1/2). We observed that the linkers had a large impact on VAP^His^ size, increasing the Stokes radius by up to ∼1.1 nm (VAP^His^ WT versus sL1/2). The VAP^His^ sL1 and sL2 constructs showed intermediate sizes and, along with the WT, followed a slope steeper than that established with folded globular standards, which is consistent with an increased hydrodynamic volume resulting from unstructured regions. Next, we conducted limited proteolysis experiments on VAP-A^His^ constructs using subtilisin, which preferentially cleaves flexible loop regions, and analyzed the resulting fragments over time by SDS-PAGE and mass spectrometry (Fig. 1c‒e). VAP-A^His^ WT was rapidly digested (t1/2= 1.13 min) upon subtilisin addition, whereas VAP-A^His^ sL1/2 was much more resistant (t_1/2_= 16.9 min) (Fig. 1c,e). As determined by mass spectrometry, two major fragments derived from VAP-A^His^ WT resulted from cleavages in L1 and L2 regions (Fig. 1d). By contrast, cleavages of VAP-A^His^ sL1/2 occurred in the coiled-coil region, consistent with the slower proteolysis of this construct. We also noticed a higher sensitivity to subtilisin of VAP-A^His^ sL2 compared to sL1 (Fig. 1e), likely because L1 is longer than L2, and thus an easier target for the protease. Taken together, these data demonstrate that VAP-A linkers are intrinsically disordered.

### The linkers of VAP are responsible for its flexibility but do not impact FFAT binding in vitro

Next, we purified and reconstituted the transmembrane (full-length) forms of VAP-A WT and sL1/2 (VAP-A^FL^ WT and sL1/2) in liposomes. VAP-A^FL^ constructs in detergent micelles were first mixed with solubilized lipids (phosphatidylcholine [PC] and phosphatidylserine [PS], at 95:5 mol/mol), and then the detergent was removed. Thereafter, reconstituted proteoliposomes were visualized by cryo-EM (Fig. 1f,g). Electron densities of proteins extending from the membrane were visible as dark dots corresponding to the MSP domain of VAP-A. As previously described, MSP domains extended at variable distances from the membrane due to the flexibility of VAP-A^FL^ WT (Fig. 1f‒h) (Mora et al., 2021), as predicted (Fig. S1a,b). Unlike for VAP-A^FL^ WT, MSP domains of the sL1/2 version were clearly resolved from vesicle surfaces (Fig. 1f, right), forming a continuous band of electron density evenly distant from the bilayer, suggesting that the proteins are rigid and adopt a similar orientation with respect to the plane of the membrane. Accordingly, VAP-A^FL^ sL1/2 exhibited more regular lengths extending from vesicle membranes (7.3 ± 1.5 nm) as compared to WT (10 ± 2 nm) (Fig. 1h). Strikingly, in membrane regions of high protein density forming ribbon-like fragments, VAP-A^FL^ sL1/2 showed a highly ordered organization, allowing us to observe sharp dark spots aligned parallel to the membrane (Fig. 1g, right). The quantification showed that changes in VAP-A^FL^ sL1/2 surface density (vesicles vs ribbons) only moderately affected its length, indicating a poorly flexible molecule (Fig. 1h).

To test whether removing VAP-A flexible linkers impacted its binding to FFAT-containing proteins, we mixed purified N-PH-FFAT, which mediates the tethering function of OSBP, with liposomes containing PI(4)P, and with VAP-A^His^ WT or mutants. We recovered the liposomes and associated proteins by flotation assay on a sucrose gradient. All VAP-A linker mutants showed the same ability to bind N-PH-FFAT binding as WT (Fig. S1c). Moreover, an excess of FFAT peptide competed with N-PH-FFAT binding to both VAP-A^His^ WT and mutants, indicating that VAP-A linkers do not directly influence MSP‒FFAT interaction.

Next, we performed cryo-EM imaging of reconstituted membrane contacts in which VAP-A^FL^ WT or sL1/2 proteoliposomes were mixed with lipid nanotubes doped with 4 mol% PI(4)P and decorated with N-PH-FFAT (Fig. S1d). Reminiscent to what we observed with VAP-A^FL^ WT (Mora et al., 2021), vesicles were flattened along the lipid nanotubes upon contact formation with VAP-A^FL^ sL1/2, and we clearly identified protein densities between the apposed membranes, indicating the formation of tethers. Then, we measured PI(4)P transfer by OSBP from Golgi-like liposomes (containing 4 mol% PI(4)P and Rhodamine lipid [Rho-PE]) to proteoliposomes containing either VAP-A^FL^ WT or sL1/2, using a specific PI(4)P probe (PH^FAPP^) labeled with an NBD molecule, as described previously (Mesmin et al., 2013). As shown in Fig. S1e, VAP-A^FL^ sL1/2 facilitated PI(4)P transfer between membranes at transport rates comparable to that obtained with VAP-A^FL^ WT (1.9±0.2 versus 1.7±0.1 PI4P/min/OSBP with VAP-A^FL^ sL1/2 and WT, respectively).

Overall, these results suggest that the two linkers of VAP-A are IDRs that allow VAP-A to adopt a range of conformations. However, altering VAP-A flexibility by removing its linkers does not affect its ability to interact with FFAT-containing proteins.

### VAP-A linkers condition its engagement to different MCSs

To assess the role of VAP-A flexible linkers in cells, we first established a VAP-A knockout (KO) RPE1 cell line by CRISPR/Cas9 (Fig. S2a,b, see Methods). We reasoned that VAP-A KO cells should help performing rescue experiments by avoiding potential heterodimerization between expressed VAP-A linker mutants and endogenous protein. In VAP-A KO cells, transiently expressed VAP-A WT tagged with the green fluorescent protein EGFP decorated the entire ER network, as expected (Fig. 2a). We noticed that EGFP-VAP-A WT also concentrated slightly in discrete ER subdomains, which may correspond to MCSs with other organelles, including the TGN (Fig. 2b,c), in agreement with previous observations (Mesmin et al., 2013; Wyles and Ridgway, 2004). In contrast to VAP-A WT, the subcellular localization of EGFP-VAP-A sL1/2 was dramatically shifted to large punctate structures dispersed throughout the cytoplasm (Fig. 2a,b). These structures overlapped with a mitochondria marker, suggesting massive recruitment of EGFP-VAP-A sL1/2 to ER‒Mito MCSs, at the expense of ER‒Golgi MCSs (Fig. 2c). However, when EGFP-VAP-A sL1/2 was further modified to prevent FFAT-motif binding (KM>DD mutation), it was no longer restricted to the vicinity of mitochondria, but was found throughout the ER network (Fig. 2a). This distribution resembled that of EGFP-VAP-A^KM>DD^, thereby suggesting that EGFP-VAP-A sL1/2 location near mitochondria depends on FFAT-containing interactors. We also verified that the massive shift driven by the sL1/2 mutation was not due to a regulatory defect: VAP-A linkers have putative Ser/Thr phosphorylation sites, the role of which, however, is not known (Cockcroft and Lev, 2020). Nevertheless, their mutation to Ala showed no effect on EGFP-VAP-A localization (Fig. S2c).

**Fig. 2.**
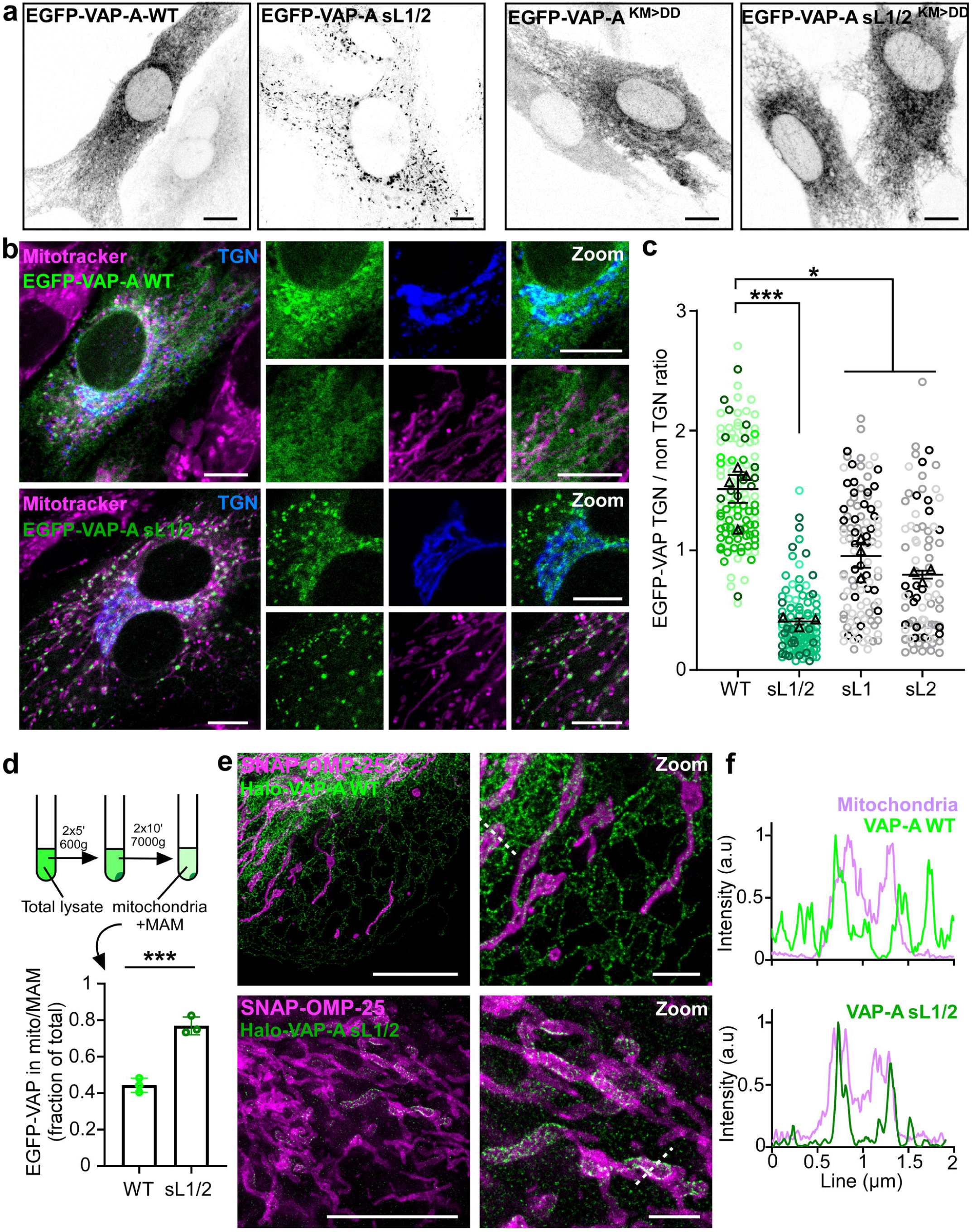
VAP-A linker mutant sL1/2 localizes mainly at contacts between the ER and mitochondria. **a**, Confocal images of RPE1 VAP-A KO cells transfected with EGFP-VAP-A WT or sL1/2 showing differential localization between the two constructs (left panels). FFAT-binding mutant EGFP-VAP-A^KM>DD^ shows no redistribution upon linker removal (right panels), suggesting that EGFP-VAP-A sL1/2 specific localization depends on FFAT-containing interactors. **b**, Cells transfected as in **a** were then stained with Mitotracker red (magenta), fixed, permeabilized and processed for immunofluorescence to assess the localization of the TGN (anti-TGN46 followed by staining with Alexa Fluor-conjugated secondary antibodies; blue). Confocal images show that EGFP-VAP-A WT decorates the entire ER and is also enriched at ER‒Golgi MCS, whereas EGFP-VAP-A sL1/2 is mainly located in punctate regions overlapping with the mitochondria marker, suggesting ER‒Mito MCSs. **c**, Superplot showing the quantification of EGFP-VAP-A constructs enrichment at the TGN area (ER-Golgi MCS). 80‒120 cells were analyzed from 3 to 4 independent experiments. Horizontal bars are mean ± SEM. **d**, Fraction of EGFP-VAP-A retrieved in purified mitochondria extract, including MAM, from cell lysate. The scheme describes the mitochondria isolation process from RPE1 VAP-A KO cell lines stably expressing EGFP-VAP-A WT or sL1/2. The graph represents mean of 3 independent experiments ± SEM. **e,** STED microscopy images of RPE-1 cells KO for VAP-A, co-transfected with Halo-VAP-A WT or sL1/2 (green) together with the mitochondria marker SNAP-OMP25 (magenta), and labelled with STED-compatible dyes before fixation. The bottom images show the recruitment of Halo-VAP-A sL1/2 around mitochondria. **f**, Fluorescence intensity of the two markers along the dashed lines from the zoomed images in **e**. Scale bars: 10 µm; 2 µm in zoomed views.

We performed subcellular organelle fractionation from VAP-A KO RPE1 cells stably expressing EGFP-VAP-A WT or sL1/2 in order to isolate mitochondria and mitochondria-associated membranes (MAM) from other organelles (Wieckowski et al., 2009). We then measured the green fluorescence of this fraction compared to that of the total lysate (Fig. 2d). Whereas ca. 40% of the signal was recovered in the Mito/MAM fraction of cells expressing EGFP-VAP-A WT, the signal almost doubled when using EGFP-VAP-A sL1/2 expressing cells. Stimulated emission depletion (STED) imaging further indicated that VAP-A sL1/2 tagged at its N-terminus with a Halo-tag was specifically recruited to regions closely apposed to outer mitochondrial membranes (OMM) (marked by SNAP-OMP25), whereas Halo-VAP-A WT largely decorated ER tubules (Fig. 2e,f and S2d). Together, these experiments provide quantitative and qualitative evidence that VAP-A sL1/2 populates ER‒Mito MCSs to a very large extent.

Compared to VAP-A sL1/2, the sL1 and sL2 mutants may retain some degree of conformational freedom, which should impact their subcellular distribution. As expected, the EGFP-VAP-A sL1 and sL2 constructs decorated the entire ER but, unlike the WT version, they concentrated at the vicinity of mitochondria and were weakly recruited at ER‒Golgi MCS (Fig. 2c and S2e). We also observed that EGFP-VAP-A sL1/2 localization at ER‒Mito MCS was attenuated when introduced into RPE1 WT cells instead of VAP-A KO cells, or co-transfected with mCherry-VAP-A WT (Fig. S2f,g). These specific contexts presumably promote the formation of heterodimers that are partially flexible.

Taken together, these results suggest that the flexibility of VAP-A provided by its linkers is necessary for its engagement in various MCSs including ER‒Golgi MCS.

### VAP-A linkers support lipid exchanges driven by OSBP and CERT at ER‒Golgi MCS

Several LTPs containing a FFAT motif, such as the cholesterol/PI(4)P exchanger OSBP and the ceramide-transfer protein CERT, are recruited by VAPs at ER‒Golgi MCS (Kawano et al., 2006; Wyles et al., 2002). These LTPs encompass a PH domain specific for PI(4)P, helping them to target the TGN (Hanada et al., 2003; Levine and Munro, 2002). As a result, their lipid transfer activities are tightly coupled to PI(4)P turnover (Capasso et al., 2017; Mesmin et al., 2017). Evidence suggests that these VAP-dependent lipid fluxes support TGN membrane remodeling and maintain TGN-to-PM vesicular traffic (Mesmin et al., 2019; Wakana et al., 2015). To test whether VAP-A flexibility is required for functional tethering at ER‒Golgi MCS, we first assessed the subcellular localization of endogenous OSBP by immunofluorescence in WT versus VAP-A KO cells, and in VAP-A KO cells stably expressing either EGFP-VAP-A WT or EGFP-VAP-A sL1/2 (Fig. 3a). OSBP was found predominantly cytosolic in WT cells, as expected, due to its PI(4)P exchange activity, which facilitates PI(4)P hydrolysis by Sac1 at the ER, and thus weakens its attachment to the TGN (Mesmin et al., 2013). However, knocking-out VAP-A largely shifted OSBP from the cytosol to the TGN, suggesting a higher PI(4)P level at the TGN resulting from a decrease in OSBP and Sac1 activities (Fig. 3a,b). Consistently, this effect could be mimicked by a 30-min treatment with 50 nM SWG, a recently described OSBP inhibitor (Fig. 3b and S3a) (Péresse et al., 2020). In this case, OSBP recruitment to the TGN was even stronger than in VAP-A KO cells, which presumably retain partial OSBP activity due to the presence of VAP-B (Fig. S2a), which is known to also interact with OSBP. In VAP-A rescue experiments, the cytosolic localization of OSBP was completely restored upon EGFP-VAP-A WT expression, but not with the sL1/2 construct (Fig. 3a,b), thereby suggesting that this mutant does not facilitate OSBP-catalyzed lipid exchange.

**Fig. 3.**
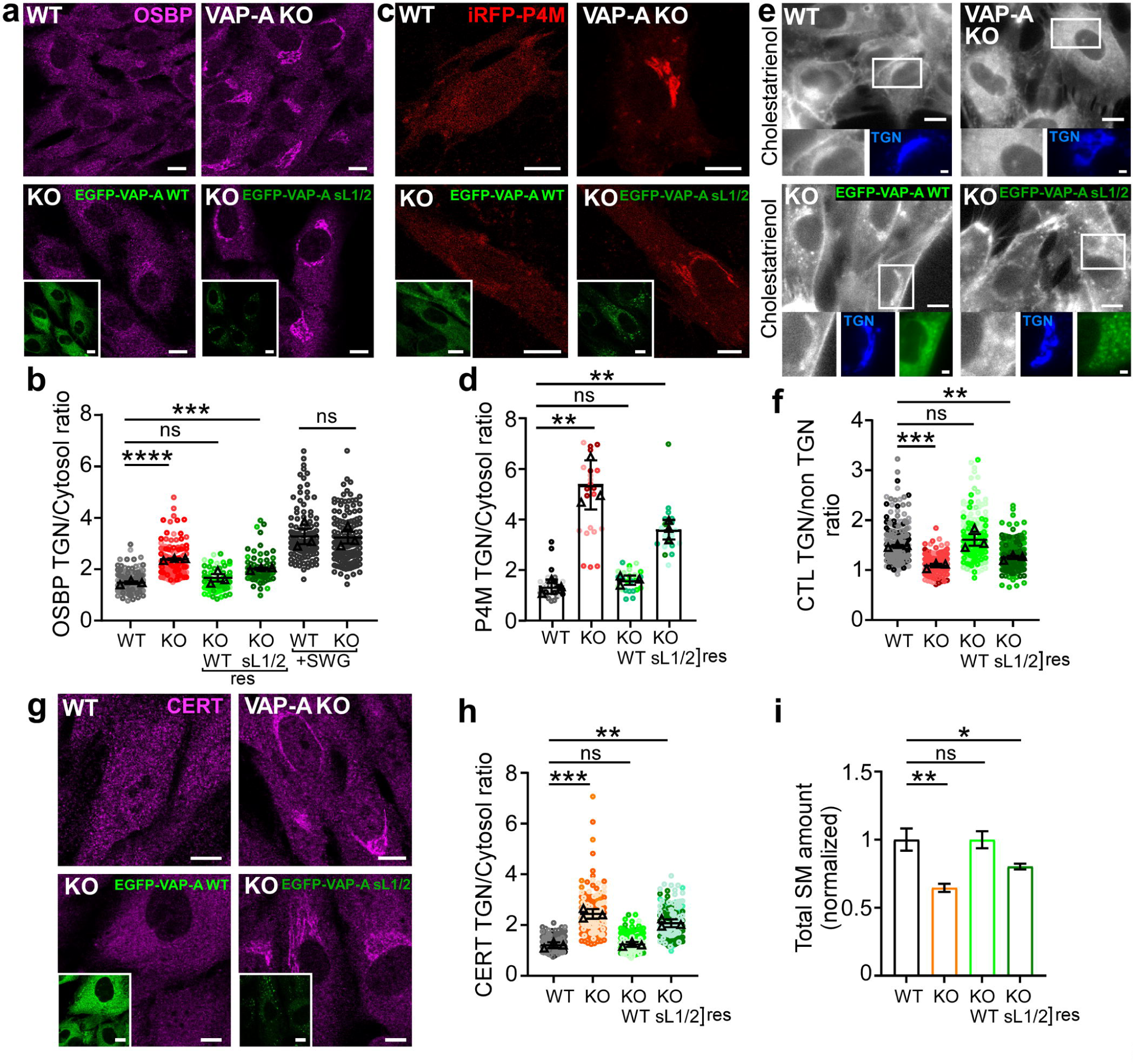
VAP-A sL1/2 does not promote OSBP and CERT-mediate lipid transport. **a**, Confocal images of endogenous OSBP visualized by immunofluorescence staining in RPE1 cells WT or KO for VAP-A. When indicated, VAP-A KO cells were stably expressing EGFP-VAP-A WT or sL1/2. OSBP is strongly recruited to the perinuclear region in KO cells or when rescued with EGFP-VAP-A sL1/2. **b**, Superplot quantifying OSBP recruitment to the TGN (res: rescue). Horizontal lines are means ± SEM from 3 independent experiments (∼150 cells analyzed). **c**, Confocal images of the PI(4)P probe iRFP-P4M transiently transfected in cells from the same conditions as in **a**. **d**, Quantification of PI(4)P probe recruitment at the perinuclear region (∼30 cells analyzed from 3 independent experiments). Lines are means ± SEM. **e**, Widefield images of RPE1 cells as in **a**, transfected with the TGN marker TagBFP-βGalT1, and pulse-chased with Cholestatrienol (CTL). Note the weak CTL labeling of the TGN in cells KO for VAP-A, or rescued with VAP-A sL1/2. **f**, Superplot showing Cholestatrienol accumulation at the TGN (n=200‒300 cells analyzed, with mean of 3 independent experiments ± SEM). **g**, Same as in **a**, except that endogenous CERT is visualized by immunofluorescence staining. **h**, Superplot showing the CERT recruitment at perinuclear area. Data are mean of 3 independent experiments ± SEM (200‒220 were cells analyzed per condition). **i**, Quantification of total SM amount by LC-MS/MS from RPE1 cells WT, KO or rescued with VAP-A WT or sL1/2. Peak areas were normalized to the control condition. Data are mean of 4 independent experiments ± SEM. Scale bars: 10 µm; 2 µm in zoomed views.

OSBP contributes significantly to intracellular cholesterol transport by consuming massive amounts of PI(4)P (Mesmin et al., 2017). We tested whether both PI(4)P turnover and cholesterol distribution were affected when replacing VAP-A WT with sL1/2. By confocal imaging, we observed that the PI(4)P probe iRFP-P4M was strongly recruited to the TGN when transfected into cells deficient for VAP-A expression (Fig. 3c,d), indicating a much larger PI(4)P pool compared with WT cells, in agreement with previous reports (Dong et al., 2016; Venditti et al., 2019a). The P4M signal at the TGN decreased sharply when cells were rescued with EGFP-VAP-A WT, but was maintained high with EGFP-VAP-A sL1/2, demonstrating a critical role of VAP-A flexible linkers in PI(4)P turnover at the TGN. To assess cholesterol distribution, we used UV-sensitive microscopy to image cells labeled with an intrinsically fluorescent analog of cholesterol, cholestatrienol (CTL, see Methods). CTL was abundant in perinuclear regions positive for the TGN marker βGalT1 in VAP-A WT cells; however, no significant CTL enrichment at the TGN could be detected in VAP-A KO cells or when rescued by EGFP-VAP-A sL1/2 (Fig. 3e,f). As an alternative method for measuring cholesterol distribution, we used the D4H probe derived from the bacterial toxin perfringolysin O (Maekawa and Fairn, 2015), which binds plasma membrane (PM) when cholesterol concentration exceeds 20 mol% of PM lipids. We previously reported that treating cells with SWG triggered D4H movement from PM to endo-lysosmal compartments, indicating that cholesterol levels dropped at PM upon OSBP inhibition (Péresse et al., 2020). We observed a similar decrease in D4H-mCherry staining of the PM in VAP-A KO cells compared to WT cells (Fig. S3b,c). In addition, this effect could not be rescued by EGFP-VAP-A sL1/2 expression.

These data suggest that the flexibility of VAP-A is an important parameter for its function as a receptor of OSBP at ER‒Golgi MCSs. To further test OSBP/VAP-A interaction, we monitored VAP-A KO cells co-expressing OSBP-mCherry and EGFP-VAP-A WT or sL1/2 by live cell imaging upon SWG treatment (Fig. S3d,e). In response to PI(4)P buildup at the TGN triggered by SWG, OSBP-mCherry and EGFP-VAP-A WT concentrated rapidly to ER‒Golgi MCS (Péresse et al., 2020), whereas no translocation of EGFP-VAP-A sL1/2 was observed, thereby indicating the lack of VAP-A sL1/2 engagement with OSBP.

Replacing endogenous VAP-A by EGFP-VAP-A sL1/2 might also impair CERT activity, as this LTP carries similar membrane targeting determinants to OSBP (Kawano et al., 2006; Levine and Munro, 2002). CERT transports newly synthesized ceramide from the ER to the TGN, where it is converted to sphingomyelin (SM) by the action of SM synthase 1 (Hanada et al., 2003). To test the requirement of VAP-A flexible linkers on CERT function at ER‒Golgi MCS, we first evaluated its subcellular localization. Staining WT cells with an antibody that recognizes endogenous CERT showed mainly a cytoplasmic signal (Fig. 3g), reminiscent to OSBP distribution, as previously described (Perry and Ridgway, 2006). By contrast, CERT showed strong perinuclear accumulation in the absence of VAP-A, which could be rescued by WT EGFP-VAP-A but not by EGFP-VAP-A sL1/2 expression (Fig. 3g,h). Interestingly, CERT relocation coincided with a significant decrease in cellular SM amounts, as determined by mass spectrometry analyses (Fig. 3i), suggesting that in such conditions, CERT was mostly bound to the TGN membrane experiencing PI(4)P buildup, as shown in Fig. 3c, but likely not engaged in functional ER‒Golgi MCS.

Taken together, these data indicate a key role for VAP-A flexible linkers in functional engagement of VAP-A with OSBP and CERT at ER‒Golgi MCSs

### Loss of VAP-A disrupts mitochondria morphology and fusion capacity

Since the less flexible VAP-A sL1/2 shows a preference for ER‒Mito MCSs, we sought to examine the role of VAP-A in these MCSs. Evidence indicates that VAP-A and VAP-B interact at ER‒Mito MCSs with several LTPs, including the OMM-anchored protein PTPIP51 (Stoica et al., 2014; Vos et al., 2012), which transfers the lipid phosphatidic acid (PA) between membranes (Yeo et al., 2021), and the channel-like proteins VPS13A and D (Leonzino et al., 2021; Li et al., 2020). VPS13A was shown to bind and transfer several phospholipids between membranes, including PA (Kumar et al., 2018). Considering that supplying PA to mitochondria supports the synthesis of cardiolipin (CL), a key lipid for the function and dynamics of mitochondria (Kameoka et al., 2018), we reasoned that VAP-A might contribute to mitochondrial morphology and dynamics. To test this hypothesis, we transfected WT or VAP-A KO RPE1 cells with a general ER marker (Sec61β-Halo) along with the OMM protein SNAP-OMP25, and performed staining with STED-compatible membrane-permeable dyes. Super-resolution imaging of WT cells showed long tubular mitochondria extending throughout the cytoplasm and making multiple contacts with the ER (Fig. 4a). By contrast, mitochondria in VAP-A KO cells were mostly fragmented and showed enlarged spherical shapes. Yet, ER contacts with round mitochondria could be detected, indicating that VAP-A is not mandatory for the formation of ER‒Mito MCS.

**Fig. 4.**
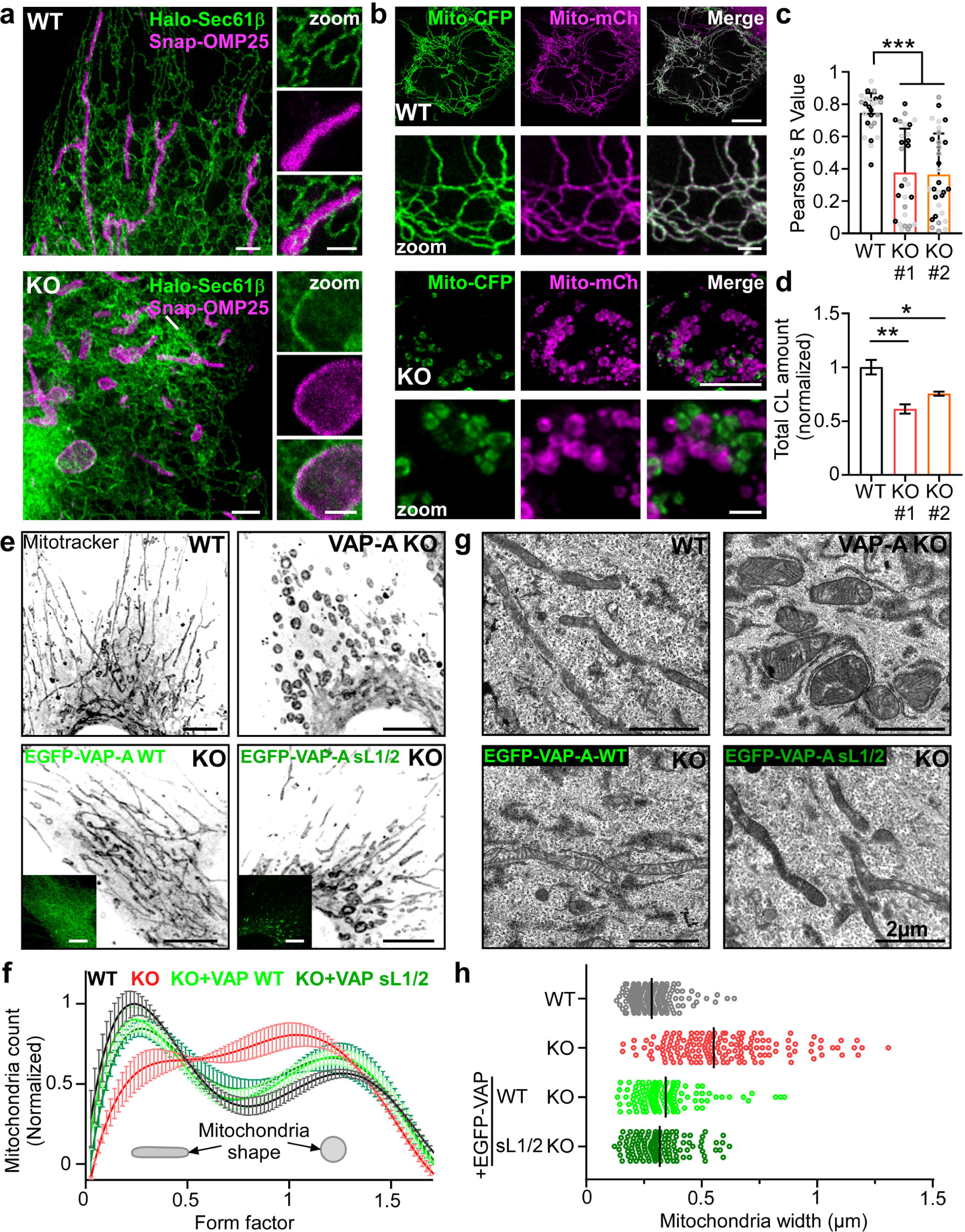
VAP-A plays a key role in mitochondrial morphology and fusion, regardless of its flexibility. **a**, Representative STED images of WT or VAP-A KO RPE1 cells expressing Halo-Sec61β (ER marker, green) and SNAP-OMP25 (magenta). Right, magnification of contact regions between the ER and mitochondria. **b**, PEG fusion assay of cells expressing mitochondrial markers in fusion with CFP or mCherry. WT RPE1 hybrid cells show extensive mitochondrial fusion (upper panels), whereas VAP-A KO hybrid cells contain mainly unfused round mitochondria (lower panels). **c**, Pearson’s coefficient colocalization measurements between Mito-CFP and Mito-mCherry. Data are mean of 3 independent experiments ± SEM (n= ∼30 fusion events analyzed). **d**, Quantification of total CL amount by LC-MS/MS of RPE1 cells WT or KO for VAP-A. Peak areas were normalized to the control condition (data are mean of 4 independent experiments ± SEM). **e**, Confocal images of WT, VAP-A KO, or VAP-A KO cells stably expressing EGFP-VAP-A WT or sL1/2, stained with Mitotracker red. **f**, Automated quantification of mitochondrial morphology (form factor) distribution. n= ∼9000 mitochondria analyzed. **g**, Thin-section EM of cells from the same conditions as in **e**. Rescues with VAP-A WT or sL1/2 reverse the mitochondrial morphology defects observed in VAP-A KO cells. **h**, Measurement of mitochondrial width from EM images. One point corresponds to one mitochondrion. Scale bars: 10 µm; 2 µm in zoomed views.

Changes in mitochondrial shape often result from unbalanced organelle fusion and fission events (Giacomello et al., 2020), but also occur during mitophagy. To test these possibilities, we first assessed the distribution of the autophagic marker LC3, which is either cytosolic in the absence of autophagy, or is recruited to forming autophagosomes. By expressing and imaging EGFP-LC3 in WT and VAP-A KO cells, we observed a similar and mostly diffuse distribution of EGFP-LC3 (Fig. S4a), distinct from that obtained upon CCCP-induced mitophagy, thereby excluding that the lack of VAP-A is associated to mitophagy. Next, we used live-cell confocal microscopy to determine whether spherical mitochondria in VAP-A KO cells resulted from aberrant mitochondrial membrane dynamics. In WT cells, frequent events of fusion and fission of mitochondria stained with MitoTracker Red were observed (Fig. S4b). By contrast, although fission occurred, no mitochondrial fusion events could be detected in VAP-A KO cells during the time course of the experiment (Fig. S4b, Movie S1). To better quantify mitochondrial fusion activity, we used polyethylene glycol (PEG) to generate hybrids between cells differentially transfected with either mitochondria-targeted CFP (Mito-CFP) or mCherry (Mito-mCherry). When WT cells were examined 16h after PEG treatment, all cell hybrids showed co-localization of fluorescence signals, indicating mitochondrial fusion (Fig. 4b,). By contrast, a significant fraction of VAP-A KO cell hybrids contained two populations of single-color mitochondria (Fig. 4b, bottom, and Fig. 4c). Thus, cells lacking VAP-A showed reduced ability to drive mitochondrial fusion.

As the phospholipids PA and CL have important roles in mitochondrial fusion, we suspected that altered fusion in VAP-A KO cells might correlate with aberrant lipid transfer and metabolism. To verify this hypothesis, we quantified total CL amounts in WT and in two independent VAP-A KO RPE1 cell lines by ultrahigh-performance liquid chromatography coupled to mass spectrometry. Compared to WT cells, CL levels decreased massively, by 43% and 25%, depending on the VAP-A-deficient cell line (Fig. 4d). PA could not be detected by lipidomic analysis, most likely due to its highly transient nature in RPE1 cells.

Overall, these experiments indicate that VAP-A contributes to the regulation of mitochondrial fusion and thus maintains mitochondrial morphology. This presumably reflects the role of VAP-A in controlling LTP-dependent PA transfer at ER‒Mito MCS, which eventually leads to CL synthesis.

### The flexibility of VAP-A is not required for its function at ER‒Mito MCS

In a next step, we investigated whether the less flexible VAP-A sL1/2 construct was functional at ER‒Mito MCS. Given that mitochondria shape was clearly different between WT and VAP-A KO cells (i.e., branched tubules versus spheres, Fig. 4a), we measured these changes using geometric parameters for precise rescue analysis. We stained mitochondria with a red fluorescent marker and performed confocal imaging of WT, VAP-A KO, and VAP-A KO cell lines stably expressing either EGFP-VAP-A WT or EGFP-VAP-A sL1/2 (Fig. 4e). As aforementioned, compared to WT cells, mitochondria were predominantly spherical in VAP-A KO cells. Strikingly, both EGFP-VAP-A WT and EGFP-VAP-A sL1/2 reversed this phenotype by restoring the population of mitochondria with a more elongated and branched shape at the expense of the spherical population (Fig. 4f, see Methods). Electron microscopy analysis confirmed these findings (Fig. 4g,h), indicating that a more rigid form of VAP-A is able to maintain mitochondrial morphology. We also compared the impact of VAP-A KO and rescues with either EGFP-VAP-A WT or sL1/2 on the levels of various CL species by lipidomics analysis. The most significant decreases upon VAP-A KO concerned CL species with fatty acid composition of low molecular weights (Fig. S4c). Their levels were substantially rescued with either VAP-A WT or sL1/2 expression.

### VAP-A interactors VPS13A and PTPIP51 maintain mitochondrial CL levels and morphology

We suspected that VPS13A and PTPIP51, which enable PA transfer between membranes, may contribute to the VAP-A dependent phenotype. We used CRISPR/Cas9-mediated gene knockout to abolish cellular expression of VAP-A, VPS13A and PTPIP51 individually or in combination (Fig. 5a, S5a,b). We then performed lipidomics analysis to measure CL amounts in the corresponding cell lines. In good agreement with a previous study showing a reduced CL level in cells silenced for PTPIP51 (Yeo et al., 2021), we observed that not only PTPIP51 KO, but also VPS13A KO, resulted in a decline of CL (by 30% and 36%, respectively), compared to WT (Fig. 5b). Of note, VPS13A/PTPIP51 double KO (DKO) produced a stronger effect, reminiscent to that obtained upon VAP-A KO (i.e., a decrease in CL of ∼45%), whereas no further drop in CL levels was measured in VAP-A/VPS13A/PTPIP51 triple KO (TKO) cells. VPS13A and PTPIP51 might thus form two pathways that converge on VAP-A. We next analyzed the phenotypic defects associated with mitochondria in these different RPE1 KO cell lines by electron microscopy. As expected, mitochondria showed inflated spherical shapes in cells knocked out for VAP-A alone or in combination with VPS13A and/or PTIPI51 (Fig. 5c,d). However, a milder phenotype was observed in VPS13A KO and PTPIP51 KO cells. In addition, since VAP-A and VAP-B have similar abundance in RPE1 cells (Fig. S5c), we also established a cell line KO for VAP-B (Fig. 5a, S5a). In contrast to what we observed with VAP-A KO, we found no significant change in CL levels in VAP-B KO cells, and no alteration in the shape of mitochondria. Overall, our results indicate that the impact of VAP-A on CL levels and on mitochondria morphology, is largely mediated by the FFAT-containing LTPs VPS13A and PTPIP51.

**Fig. 5.**
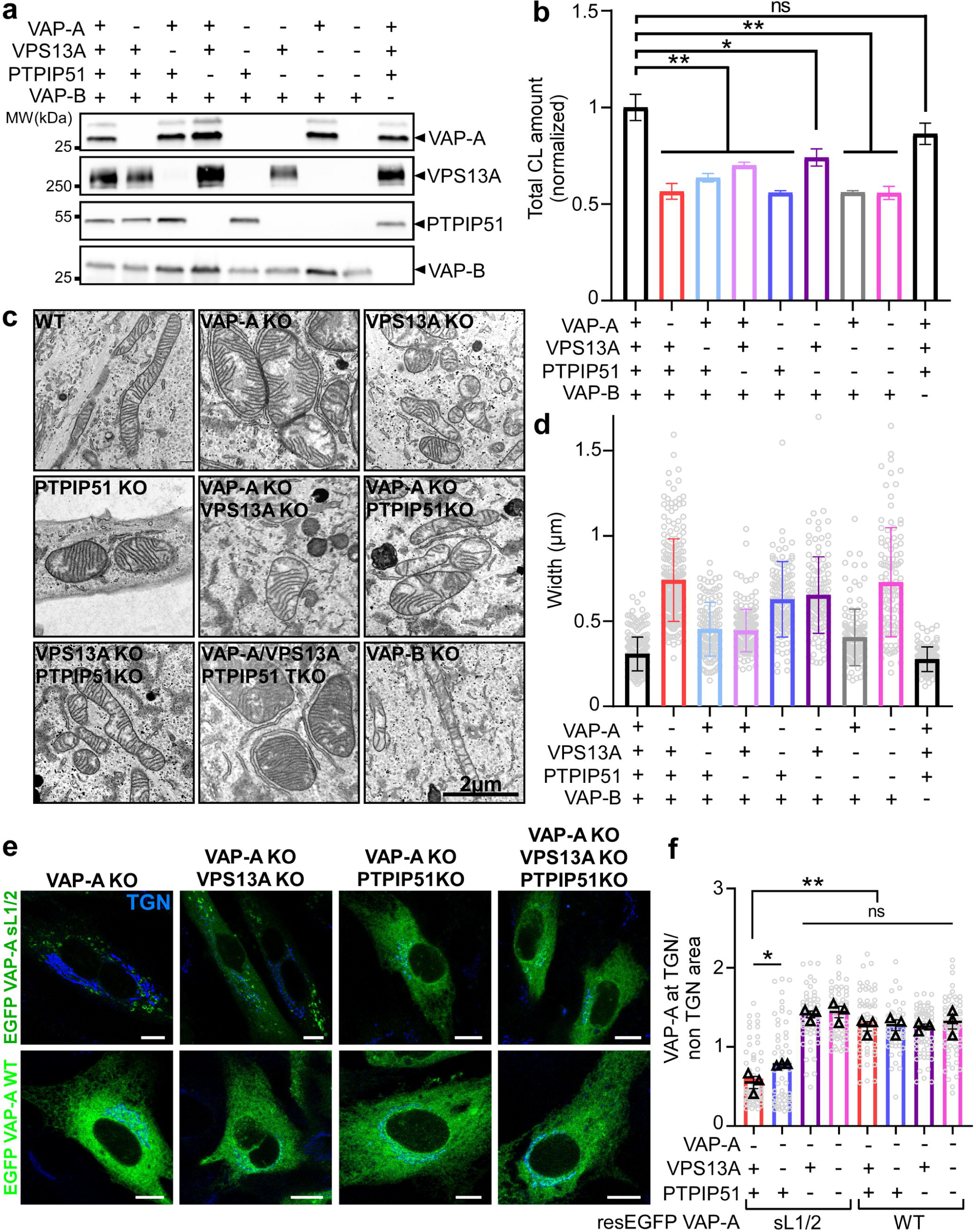
Mitochondrial shape and CL levels are affected in cells lacking VAP-A interactors VPS13A and PTPIP51. **a**, Western-blot analysis of different CRISPR/Cas9 KO RPE1 cell lines. **b**, Quantification of total CL amount by LC-MS/MS of each KO cell line. Peak areas were normalized to the control condition. Data are mean of 4 independent experiments ± SEM. **c**, Thin-section EM of the different KO cell lines emphasizing mitochondrial morphology. VPS13A and/or PTPIP51 KO cells display round and inflated mitochondria. Note that VAP-B KO cells display elongated mitochondria resembling that of WT cells. **d**, Graph showing mitochondrial width, measured for each condition. One point corresponds to one mitochondrion. **e**, Confocal imaging of cells KO for the indicated proteins and transiently transfected with EGFP-VAP-A sL1/2 or WT. The absence of VAP-A interactors, VPS13A and especially PTPIP51, results in the redistribution of EGFP-VAP-A sL1/2. **f**, Quantification of EGFP-VAP-A enrichment at the TGN region. Data are mean of 3 independent experiments ± SEM (n= ∼60 cells analyzed). Scale bars: 10 µm; 2 µm in zoomed views.

Given that VAP-A sL1/2 was mostly found at ER‒Mito MCSs (Fig. 2b), the above analysis suggested that VPS13A and PTPIP51 may account for its localization. To test this hypothesis, we monitored EGFP-VAP-A sL1/2 subcellular distribution in VAP-A KO cells lines lacking VPS13A and/or PTPIP51 expression. As shown in Fig. 5e, the sL1/2 construct was no longer restricted to the vicinity of mitochondria in VAP-A/VPS13A DKO, VAP-A/PTPIP51 DKO and VAP-A/VPS13A/PTPIP51 TKO cells. The effect was more striking in the last two conditions where EGFP-VAP-A sL1/2 relocated all over the ER, reminiscent of EGFP-VAP-A WT distribution (Fig. 5f). These results demonstrate that VPS13A and PTPIP51 are key partners of VAP-A at ER‒Mito MCSs, and the primary interactors of VAP-A sL1/2.

### Flexible linkers enable VAP-A to adapt its function to short lived-MCS but do not modulate partner preference

The experiments described so far suggest that owing to its flexibility, VAP-A could accommodate partners of different sizes and/or structures, whereas a more rigid tether would have a limited interaction range. An alternative hypothesis, however, is that VAP-A might promote membrane tethering at many MCSs, whereas VAP-A sL1/2 might be selective according to MCS structural stability, regardless of the exact interactor nature. To address how VAP-A flexible linkers determine its tethering properties, we first considered reducing the dynamics of ER‒Golgi MCSs, which are transient in nature. This can be achieved by overexpressing the PH-FFAT fragment of OSBP, which by interacting with VAP-A produces membrane bridges that persist over time (Mesmin et al., 2013; Wakana et al., 2015). As depicted in Fig. 6a, EGFP-VAP-A WT concentrated at the perinuclear area upon PH-FFAT-mCherry expression. Fluorescence recovery after photobleaching (FRAP) indicated that PH-FFAT and VAP-A exchanged very slowly from these regions, which is consistent with stable membrane tethering at ER‒Golgi MCSs (Fig. 6b, S6a). Strikingly, PH-FFAT-mCherry also induced robust relocation of EGFP-VAP-A sL1/2, which no longer decorated ER‒Mito MCS (Fig. 6a). As a control, we introduced a point mutation in PH-FFAT-mCherry (R108L) that strongly reduces its affinity for PI(4)P, but does not abrogate Golgi targeting (Levine and Munro, 1998; Mesmin et al., 2013). As expected, PH-FFAT-mCherry R108L recovery was much faster (∼3 fold) than the corresponding WT construct by FRAP (Fig. 6b), indicating that ER‒Golgi MCSs were poorly stabilized in this case. Although EGFP-VAP-A WT was efficiently relocated to the perinuclear region by PH-FFAT-mCherry R108L, EGFP-VAP-A sL1/2 was not and, instead, distributed predominantly at ER‒Mito MCSs (Fig. 6a). Colocalization quantification confirmed these differential relocations (Fig. 6c).

**Fig. 6.**
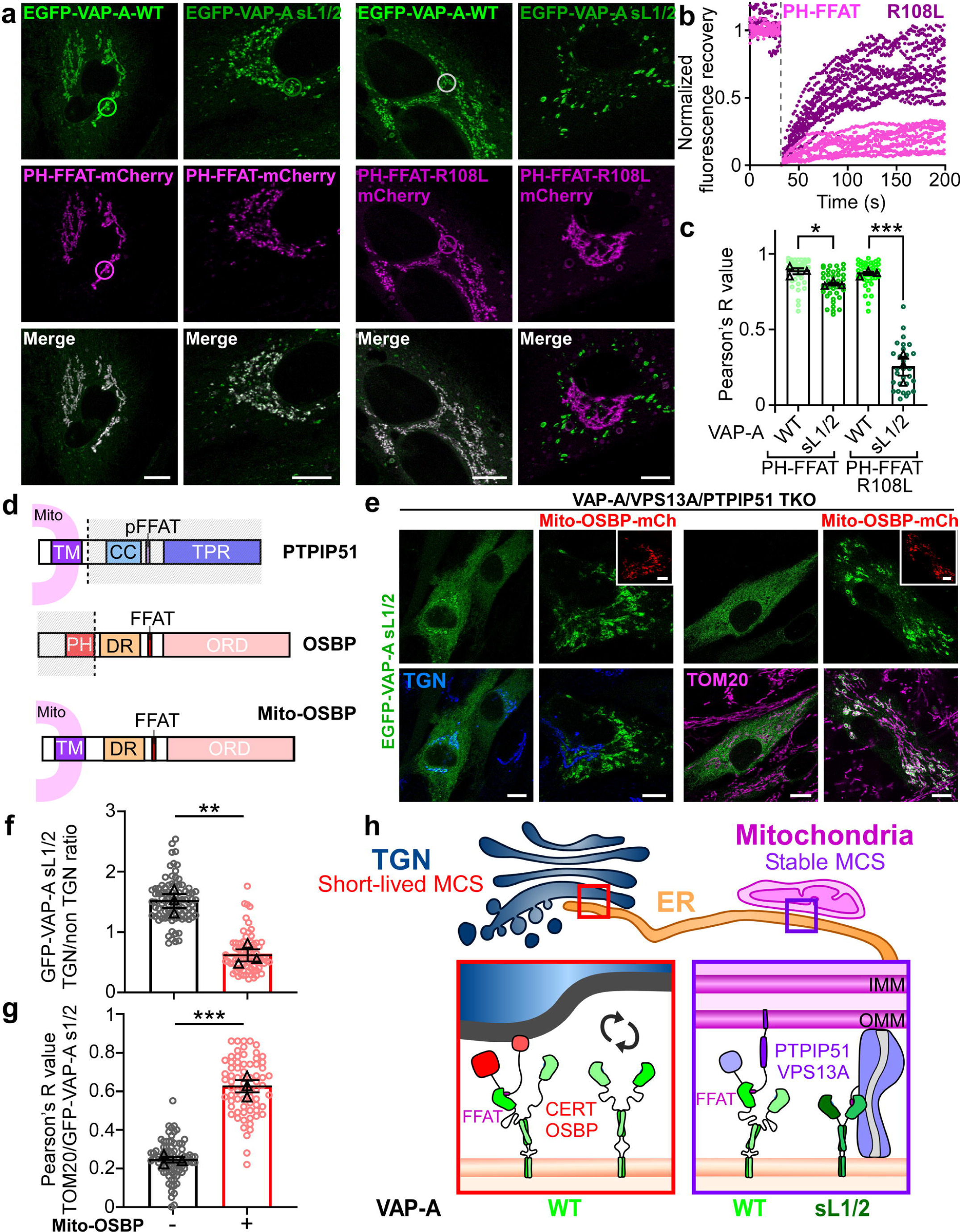
VAP-A sL1/2 shows preference for stable MCSs, regardless of its interactors. **a**, Confocal images of VAP-A KO RPE1 stably expressing EGFP-VAP-A WT or sL1/2 and further transfected with PH-FFAT-mCherry or PH-FFAT-R108L-mCherry. The R108L mutation weakens PH domain binding to PI(4)P. Both EGFP-VAP-A construct are strongly recruited to the TGN area upon PH-FFAT expression, which stabilizes ER‒Golgi MCS, but not with PH-FFAT-R108L, which recruits only EGFP-VAP-A WT. **b**, FRAP analysis of PH-FFAT-mCherry WT or R108L from cells coexpressing EGFP-VAP-A WT. 10 cells were analyzed in 2 independents experiments. **c**, Colocalization test between EGFP-VAP-A WT or sL1/2 and PH-FFAT-mCherry WT or R108L. Data represent means of 3 experiments ± SEM (∼50 cells were analyzed,). **d**, Domain organization of PTPIP51, OSBP, and Mito-OSBP, a chimeric construct in which the PH domain of OSBP is replaced by the transmembrane domain of PTPIP51. **e**, Confocal images of RPE1 KO for VAP-A, VPS13A and PTPIP51, transiently expressing EGFP-VAP-A sL1/2 and Mito-OSBP-mCherry (red, when indicated), further processed for immunofluorescence to detect either the TGN (anti-TGN46, blue) or mitochondria (anti-TOM20, magenta). Mito-OSBP expression restores EGFP-VAP-A sL1/2 localization at ER‒Mito MCS in cells lacking endogenous VAP-A interactors. **f**, Quantification of EGFP-VAP-A sL1/2 enrichment at TGN area. Data are mean ± SEM from 3 independent experiments (∼90 cells were analyzed). **g**, Pearson’s colocalization test between TOM20 and EGFP-VAP-A sl1/2 depending on Mito-OSBP-mCherry expression (mean of 3 independent experiments ± SEM; 85 cells analyzed). **h**, Model showing that flexible linkers enable VAP-A to explore a larger conformational space to capture partners, and thus adapt its function to various types of MCS including short-lived ER‒Golgi MCS, whereas a rigid VAP-A mutant is restricted to long-lived MCSs. Scale bars: 10 µm; 2 µm in zoomed views.

To further demonstrate that the lack of flexible linkers restricts the ability of VAP-A to adapt to different MCSs rather than different partners, we sought to attach a Golgi-specific VAP-A interactor to mitochondria in cells lacking the mitochondria-specific VAP-A interactors VPS13A and PTPIP51. For this, we designed a mCherry-tagged OSBP construct in which its N-terminal segment comprising the PH domain (OSBP res. 1-178) was replaced by the TM domain of PTPIP51 (res. 1-43) (Fig. 6d). Confocal imaging confirmed that the OSBP chimera (Mito-OSBP-mCherry) relocated to mitochondria in VAP-A/VPS13A/PTPIP51 TKO cells (Fig. 6e). Thereafter, EGFP-VAP-A sL1/2 subcellular distribution was assessed. Strikingly, the sL1/2 construct was no longer distributed throughout the ER, but concentrated near mitochondria in the presence of Mito-OSBP-mCherry, as measured by the strong increase in Pearson’s coefficient between EGFP-VAP-A sL1/2 and the OMM marker TOM20 (Fig. 6e‒g, S6b,c).

Taken together, the two experimental strategies shown in Fig. 6 indicate that rigid VAP-A sL1/2 preferentially concentrates in MCS with greater stability, whereas the flexible linkers of VAP-A allow its engagement in highly dynamic MCSs, notably short-lived ER‒Golgi MCS.

## DISCUSSION

Here, we investigated the functional architecture of VAP-A, a receptor that contributes significantly to the function of the ER, by interacting with a myriad of proteins involved in multiple functions and by bridging the ER to other organelles. We identified two linker regions in VAP-A as key elements controlling its tethering activity. These regions appear to be unstructured as judged by their proteolytic sensitivity, their behavior in gel filtration, and their behavior in MD simulations. This is consistent with the fact that the length of VAP-A is variable according to its membrane density (Mora et al., 2021). Remarkably, cryo-EM visualization of a VAP-A construct in which both linker regions were replaced by short tracts (VAP-A sL1/2) indicated that it is more rigid than the WT form. Although VAP sL1/2 retains its intrinsic capacity to interact with FFAT-containing proteins, a major observation was its massive relocalization to ER‒Mito MCS at the expense of ER‒Golgi MCS. As a result, the function of VAP-dependent LTPs OSBP and CERT in Golgi lipid traffic was impaired whereas VAP-A sL1/2 could recapitulate VAP-A WT function at ER‒Mito MCS. These include supporting CL metabolism and mitochondrial fusion capacity through interactions with PTPIP51 and VPS13A.

Many proteins enriched at MCS contain segments predicted to be IDRs (Gatta and Levine, 2017; Jamecna and Antonny, 2021); however, their roles and impact on protein dynamics remain largely unexplored. When unfolded linker regions are long enough, such as in Ist2, their primary function at MCS could be to span interorganellar gap to form a molecular bridge between membranes (D’Ambrosio et al., 2020; Kralt et al., 2015). Alternatively, short linkers may restrict the action range of catalytic domains. Such mechanism prevents the phosphatase Sac1 from hydrolyzing PI(4)P *in trans* at MCS (Zewe et al., 2018). Our recent study on OSBP showed that its long N-terminal IDR enables decreasing protein density at MCS and facilitating protein lateral movements (Jamecna et al., 2019). With the example of VAP-A, our work reveals a key role for IDRs at MCS: to enable membrane tethering at several MCSs showing different dynamics.

At first sight, one can simply consider that VAP-A sL1/2 accumulates at ER‒Mito MCS due to its smaller size compared to VAP-A WT, such as to adapt to its partners’ sizes or to a precise intermembrane distance. However, we showed that VAP-A sL1/2 relocated back at ER‒Golgi MCS when such MCS are artificially stabilized. Alternatively, positioning OSBP at mitochondria surface could promote VAP-A sL1/2 localization at ER‒Mito MCS in cells lacking VPS13A and PTPIP51. Thus, a size-mediated segregation mechanism is unlikely to occur. Akin to adhesion molecules mechanics at cell‒cell contacts (Jeppesen et al., 2001; Li et al., 2021), we propose that VAP-A IDRs behave as molecular springs that ensure its vertical flexibility to provide adaptive tethering at MCS, which would facilitate membrane‒membrane adhesion when intramembrane distance fluctuates. The model in Fig. 6h shows that owing to its flexible IDRs, VAP-A tethers membranes in any MCS contexts, including ER‒Golgi MCSs, which are highly dependent on PI(4)P turnover, and thus subjected to fluctuations, whereas VAP-A sL1/2 requires more stable contacts to efficiently interacts with partners. Of note, simulations of coarse-grained membrane apposition systems have shown that rigid membrane-anchored receptors and ligands are more prone to dissociation when membrane shape fluctuations increase (Hu et al., 2013). Apart from ER‒Golgi MCS, other MCSs also show a dependence on phosphoinositide metabolism, which may lead to rapid membrane association and dissociation events. According to their relative stiffness, tethers may be included in or excluded from such MCSs.

Our work indicates that VAP-A acts as a key player in mitochondrial fusion, suggesting a hitherto unknown physiological role for this general ER receptor. We observed that morphological changes in mitochondria in the absence of VAP-A, or of its interactors PTPIP51 and VPS13A, which transfer PA between membranes, correlated with a drastic decrease in mitochondrial CL content. Interestingly, similar mitochondrial shape alterations were also reported in Drosophila KO for VPS13D (Anding et al., 2018). Following CL synthesis in the IMM, tafazzin-mediated CL remodeling occurs, which results in polyunsaturated species (Schlame and Xu, 2020). Our lipidomic analyzes indicated that the CL species that predominantly decreased upon VAP-A KO exhibit low molecular weight and more saturated fatty acyl chains (Fig. S4c). This might suggest that neosynthesized species are primarily affected, thereby supporting the role of VAP-A in facilitating the transfer of PA, the precursor of CL, to mitochondria.

How CL promotes mitochondrial fusion has been demonstrated by several studies. Notably, CL is directly involved in membrane fusion through interactions with OPA1 (Ban et al., 2017); yet this reaction concerns mitochondrial inner membrane fusion. At the outer membrane, mitofusin-mediated fusion is stimulated by the enzyme MitoPLD that hydrolyzes CL to generate PA (Choi et al., 2006), which has presumably fusogenic properties due to its small polar head. Of note, MitoPLD downregulation also induced a mitochondrial fragmentation phenotype reminiscent of VAP-A KO. Moreover, it was reported that Mitoguardin (MIGA) proteins stabilize MitoPLD to promote mitochondrial fusion (Zhang et al., 2016). Interestingly, Klemm *et al*. recently revealed that MIGA2 is able to interact with VAP through a FFAT motif, and thus participates to ER‒Mito tethering (Freyre et al., 2019). These elements taken together place VAP-A as a central factor controlling mitochondria lipid metabolism and membrane dynamics.

Previous studies have mostly examined the role of VAP-B at ER‒Mito MCSs, rarely that of VAP-A. However, VAP-A is abundantly present in these MCSs. Surprisingly, our data show divergent implications of VAP-A and VAP-B on CL levels and mitochondrial morphology (Fig. 5b‒d), even though they have comparable expression levels. Of note, unlike VAP-B, VAP-A knockout is embryonic lethal in mice (Kabashi et al., 2013; McCune et al., 2017). Interestingly, although the MSP domains of VAP-A and VAP-B are nearly identical, their linkers and predicted coiled-coils are substantially different (41.7% sequence identity, compared to 82.2% for MSP). It is possible that variations in VAP “stem” region could contribute to interaction/localization preferences. Whether VAP-A has functions that are not shared with VAP-B is an important issue that will require further studies.

The fact that VAP-A sL1/2 restored mitochondria morphology in VAP-A KO cells suggests that the folded domains of VAP-A are sufficient to guarantee an important part of its function at ER‒Mito MCS. This highlights the molecular and functional heterogeneities between ER‒Golgi and ER‒Mito MCSs, which most likely involve changes in intermembrane distances fluctuations coupled to the dynamics of their specific components. The thickness of a single MCS type, often observed by EM in resin embedded samples or by in situ cryo-EM, shows very large variations from the molecular point of view (Hela ER‒Golgi MCS: 5‒20 nm; yeast ER‒Mito MCS: 16±7 nm) (Collado et al., 2019; Venditti et al., 2019b), as they likely represent snapshot of transient states. Determining intermembrane distance fluctuations with high spatiotemporal resolution will be helpful in the future to finely describe MCS structure and dynamics.

## ACKNOWLEDGMENTS

We thank F. Brau, N. Leroudier (IPMC, Valbonne) and J. Manzi (Institut Curie, Paris) for technical assistance; Z. Riegel, A. Lamy, L. Velut, R. Charreire, V. Lejal, C. Watine and S. Goffart (Polytech Nice Sophia engineer school) for providing useful tools and scripts; K. Schauer (IGR, Paris) for discussions; F. Bottanelli (FU Berlin), F. Alpy (IGBMC, Illkirch) and E. Honoré (IPMC, Valbonne) for plasmids, and all members of the Antonny and Lévy labs for their insights. We thank the IPMC Imaging and Cytometry facility, part of the MICA platform (GIS IBiSA), and the Cell and Tissue Imaging core facility (PICT IBiSA), Institut Curie, member of the French National Research Infrastructure France-BioImaging (ANR-10-INBS-04). This work was supported by the CNRS, the Inserm, the Agence Nationale de la Recherche (ANR-21-CE13-0021-01) (to B.M., D.L. and F.R.) and the ERC (Synergy #856404) (to B.A.). M.S. was supported by a PhD fellowship from the Université Côte d’Azur, Académie 4 (ANR-15-IDEX-01).

## AUTHOR CONTRIBUTIONS

B.M. and D.L. conceived the study. B.M., M.S. and D.L. designed experiments. M.S. performed experiments. M.D., J.B. and B.M. purified proteins. A.D.C., M.D. and D.L. performed cryo-EM imaging and analysis. S.L.C. performed EM imaging of cell samples. M.S. and S.A. performed STED imaging. A.R.D.A., L.F. and D.D. performed lipidomics and mass spectrometry measurements. R.G. performed molecular dynamics simulations. F.R. synthesized cholestatrienol. B.A. and A.P. provided advice on experimental design. B.M. and M.S. wrote the manuscript. B.M., B.A. and D.L. reviewed and edited the manuscript.

## METHODS

## RESOURCES TABLE

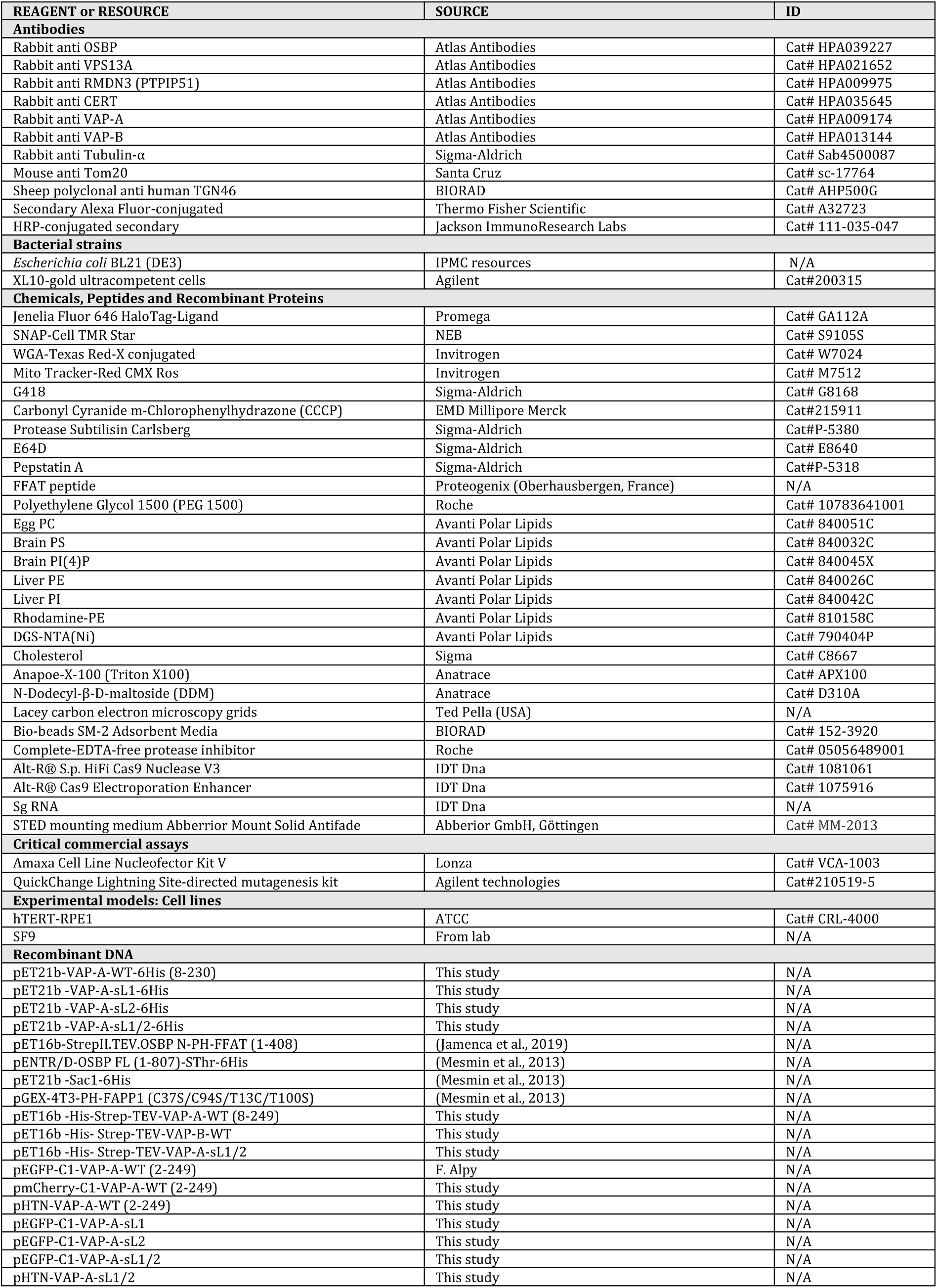
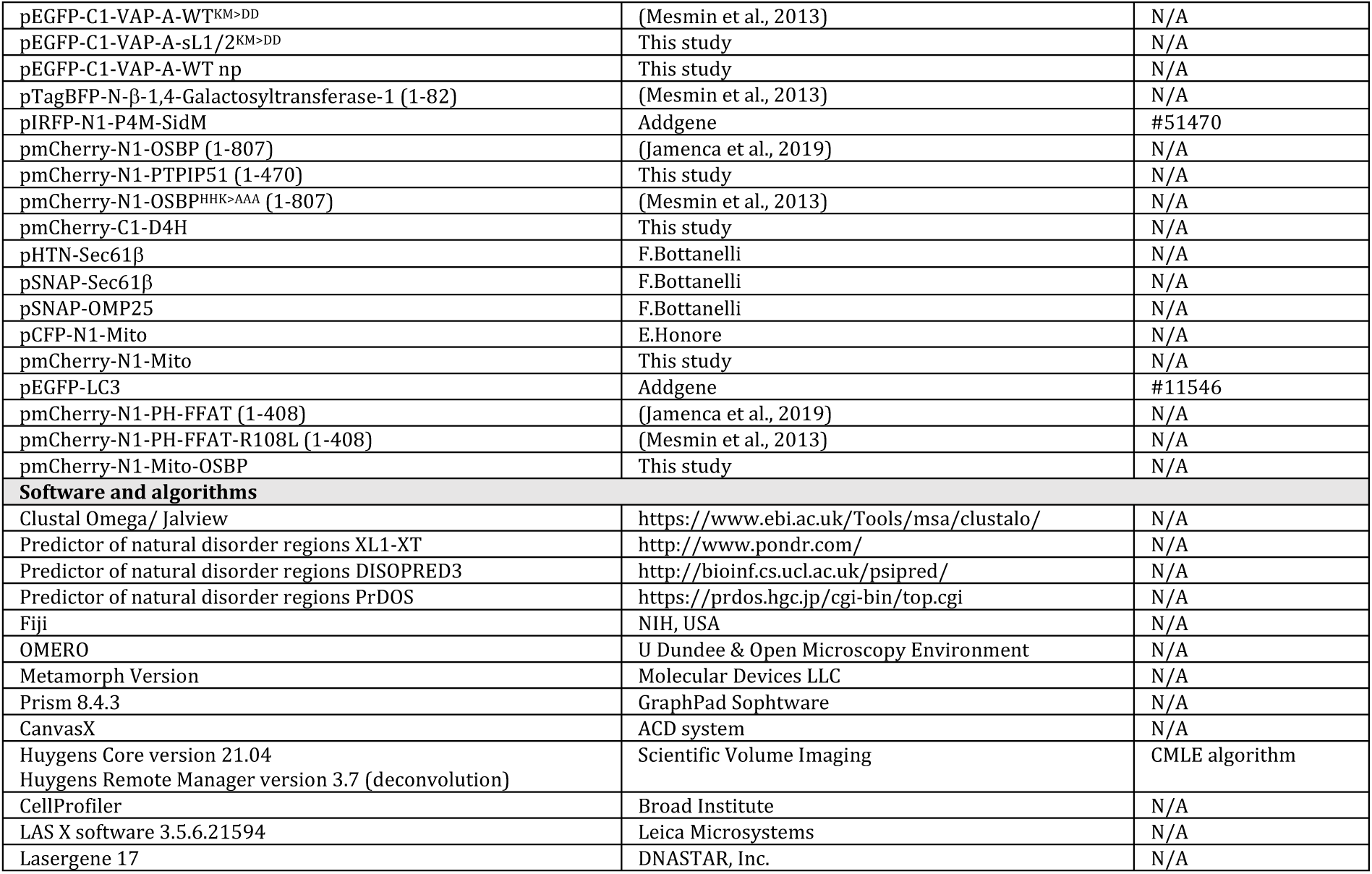

## CONTACT FOR REAGENT AND RESOURCE SHARING

Further information and requests for plasmids and reagents should be directed to and will be fulfilled by the lead contact, Bruno Mesmin.

## EXPERIMENTAL MODEL AND SUBJECT DETAILS

### Cell culture

#### hTERT-RPE1 cells

Human immortalized retinal pigmented epithelial cells (hTERT-RPE1, ATCC Cat# CRL-4000; hereafter RPE1) were used as cellular model. RPE1 are genetically stable, lack transformed phenotypes, and are commonly used to study intracellular membrane traffic and dynamics. They were cultured in DMEM/F12 (Gibco) supplemented with 10% fetal calf serum (FCS) and 1% antibiotics (Zell Shield) at 37°C in a 5% CO2 humidified atmosphere.

#### Insect (Spodoptera frugiperda) SF9 cells

SF9 cells were cultured at 27°C in SF-900 II media (Gibco) supplemented with 1.5% FCS in absence of antibiotic.

## METHODS DETAILS

### Plasmids

To reduce VAP-A flexibility while preserving the rotation and dimerization properties of the protein, the flexible linkers between MSP and CC (L1; aa. 133-164) and between CC and TM domains (L2; aa. 207-230) were replaced by short Gly/Ser-rich sequences GGSGGSGG (sL1) and GSG (sL2). Mutagenesis was performed from three starting vectors: pEGFP-C1 VAP-A (FL; 1-249), pET21b+ VAP-A His6 (8-230) and pET16b His6-StrepII-TEV VAP-A (8-249) by Eurofins Genomics. The chimeric construct Mito-OSBP was cloned into a pmCherry N1 vector. For this, the transmembrane domain of PTPIP51 (aa. 1-43) was created by annealing two long primers covering the entire sequence and OSBP fragment [179-807] was amplified by a PCR reaction. Both fragments had an additional end that hybridizes each other. A second PCR was done with both templates and oligos containing NheI and BamHI restriction sites. The final insert was digested as well as the vector before ligation and transformation into XL10 Gold cells. Other constructs from this study were obtained using the restriction enzyme cloning method. Point mutations to obtain EGFP-VAP-A sL1/2 KM>DD and EGFP-VAP-A np were performed using the Quickchange lightning mutagenesis kit (Agilent). For VAP-A sL1/2 KM>DD, Lys 94 and Met 96 were replaced by Asp; for VAP-A np, Ser 145, 164, 214, 216, 219 and the Thr 215 were replaced by Ala. All plasmid constructs were sequenced, and analyzed with DNASTAR Lasergene software.

### Cell transfection

RPE1 cells were transfected by electroporation with Nucleofector solution V (Lonza) using the Amaxa Nucleofector device (X-01 program) (Lonza) for 24-48 h. RPE1 cells stably expressing EGFP-VAP were selected using G418 (500 mg/mL) (Sigma). Surviving colonies were isolated using cloning cylinders (Bel-Art). For microscopy, cells were seeded at suitable density to reach 50-90% confluence on the day of imaging on ibiTreat μ-Slides (Ibidi) or in uncoated glass coverslip for STED experiments.

### Protein expression and purification

Full length human VAP-A WT and sL1/2 expression were performed in C41 (DE3) *E. coli* strain in presence of 0.5 mM IPTG for 4 h at 30°C, and purified according to (Mora et al., 2021). VAP expressing cells were collected, lysed through a cell disruptor and centrifuged to purify membranes. Then, membranes were solubilized for 2h at 4°C with N-Dodecyl-β-D-maltoside (DDM):protein ratio of 2.5 (w/w) and then centrifuged to collect solubilized proteins. VAP-A WT and sL1/2 were purified on Strep Tactin beads (IBA superflow sepharose) in batch. Eluted proteins were incubated overnight with TEV protease to eliminate the double tag, followed by purification on a Superdex 200 (GE healthcare) size exclusion chromatography in Tris 50 mM pH 7.5, NaCl 150 mM, DDM 30 mM, 10% glycerol. Purified proteins were collected at concentration ranging from 1.5 to 2 mg/mL without any additional concentration step, flash frozen in liquid nitrogen and stored at -80°C. Full length human VAP-B was purified using the same method.

OSBP N-PH-FFAT, Sac1^His^ and VAP-A^His^ WT, sL1, sL2 and SL1/2 constructs, sub-cloned into pET-21b, were expressed in *E. coli* BL21 (DE3) and purified according to refs. (Jamecna et al., 2019; Mesmin et al., 2013). N-PH-FFAT had a N-terminal 6xHis tag-StrepII-TEV site, VAP-A^His^ constructs and Sac1^His^ (1-522) had a C-terminal 6xHis-tag replacing protein transmembrane region. After protein expression, bacteria were lysed in a cell disruptor at 1600 psi and incubated for 30 min on ice with DNAse and MgCl2 (5mM) before ultracentrifugation (125,000×g). Proteins were purified using HisPur^TM^ Cobalt Resin (Thermo Scientific). For N-PH-FFAT, protein fractions were pooled and cleaved with TEV protease overnight at 4°C. Digested proteins were purified on SourceQ HR 10/10 column (GE Healthcare) with a 0-1M NaCl gradient in 25mM Tris pH7.5 followed by a Sephacryl S200 HK16/70 column (GE Healthcare) equilibrated in 25mM Tris pH7.5, 120 mM NaCl, 2mM DTT, using an AKTA purifier system (GE Healthcare). For VAP^His^ constructs and Sac1^His^, eluted protein fractions were pooled, concentrated and further purified on a Sephacryl S200 HR/XK 16/70 column (GE Healthcare) equilibrated in 25 mM Tris pH 7.5, 120 mM NaCl, 1 mM DTT.

PH-FAPP1 in pGEX-4T3, modified by point mutagenesis (C37S, C94S, T13C, T100S) for specific NBD labeling of C13, was expressed as a GST fusion protein in *E.coli* and purified according to ref. (Mesmin et al., 2013) on Glutathione Sepharose 4B beads (GE Healthcare), cleaved by thrombin at 4°C overnight and next purified by gel filtration on a Sephacryl S300 HR XK16–70 column in 50 mM Tris, pH 7.4, 120 mM NaCl and 2 mM DTT. For NBD labelling, purified PH-FAPP1 was first passed on a NAP-10 column (GE Healthcare) to remove DTT and then mixed with a 10-fold excess of N,N’ -dimethyl-N-(iodoacetyl)-N’ -(7-nitrobenz-2-oxa-1,3-diazol-4-yl) ethylenediamine (IANBD-amide, Molecular Probes). After 90 min on ice, the reaction was stopped by adding a 10-fold excess of L-cysteine. The free probe was removed by gel filtration on a Sephacryl S200 HR XK16–70 column.

Full length (1-807) OSBP was purified from baculovirus-infected SF9 cell according to ref. (Mesmin et al., 2013). The construct was created by subcloning PCR-amplified DNA into pENTR/D-Topo vector (Invitrogen), followed by insertion of a thrombin site at the end of the OSBP sequence. The PCR fragment was then inserted by recombination into linearized BaculoDirect vector (C-terminal V5 and His-tagged), and transfected into SF9 cells. OSBP with a C-ter 6×His tag was produced in Sf9 cells, which were infected with baculovirus at MOI of 0.1 for 1h at room temperature, suspended in 500–1000 mL of Sf-900 II media supplemented with 1.5% FCS and incubated with mixing at 27°C for 72 h. Thereafter, cells were collected by centrifugation at 300×g for 10 min and pellets were stored at -20°C. Cell pellets were resuspended in lysis buffer (20 mM Tris pH 7.5, 300 mM NaCl, 20 mM imidazole, EDTA-free protease inhibitors and phosphatases inhibitors) and lysed with Dounce homogenizer. After ultracentrifugation (125,000×g), OSBP from the supernatant was adsorbed on an HisPur^TM^ Cobalt Resin (Thermo Scientific), submitted to 3 washes with lysis buffer supplemented with 800, 550, and 300 mM NaCl, respectively, and then eluted with 250 mM imidazole-containing buffer. OSBP fractions were pooled, concentrated on Amicon Ultra (cut-off 30 kDa), and submitted to thrombin cleavage for 1 h at 25°C to eliminate the His tag, then purified on a Sephacryl S300 HK16/70 column (GE Healthcare). All steps were performed at 4°C. Purified proteins were concentrated, supplemented with 10% glycerol, flash-frozen in liquid nitrogen and stored at -80°C.

### Analytical Gel filtration

Purified VAP-A^His^ constructs (100 µL, 30 µM) were applied on a Supedex 75 column and eluted at a flow rate of 0.5 mL/min in 25 mM Tris pH 7.5, 120 mM NaCl and 1 mM DTT. The column was calibrated using the following standards (MW/Stokes radius): Alcohol Dehydrogenase (150 kDa/46 Å), Bovine Serum Albumin (67 kDa/36.1 Å), Ovalbumin (43 kDa/28 Å), Carbonic Anhydrase (29 kDa/21 Å). The elution volume and Stokes radius of the standards were used to establish a first calibration curve, from which the Stokes radius of the VAP^His^ constructs were determined. Thereafter, the Stokes radius was potted as a function of MW for both protein standards and for VAP-A^His^ constructs.

### Limited proteolysis

VAP-A^His^ WT, sL1, sL2 or sL1/2 (4 µM) were mixed in 50 mM HEPES pH 7.4, 120 mM K acetate, 1 mM MgCl2 (HKM) buffer. Subtilisin (1 µg/mL) was added at time zero, and, at the indicated times the reaction was stopped by adding 0.5 µM Phenylmethylsulfonyl fluoride. The samples were analyzed by SDS PAGE with Sypro-Orange staining and by liquid chromatography coupled to a mass spectrometer equipped with a heated electrospray ionization (HESI) probe. HPLC was performed using a Dionex U3000 RSCL Instrument (ThermoFisher Scientific). Protein analysis was performed on a Syncronis C18 column at flow rate of 300 µL/min. The elution program was based on water (solvent A) and acetonitrile (solvent B) both containing 0.1% formic acid (v/v): 0 min 3% B, 25 min 60% B, 28 min 95% B. The Q-exactive *plus* spectrometer controlled by the Xcalibur software was operating in electrospray positive mode. Typical ESI conditions were as follows: electrospray voltage 4 kV; capillary temperature 320°C, probe temperature 325°C, sheath gas flow 40U and auxiliary gas 10U. The MS scan was acquired in the 400 to 1900 m/z range with the resolution set to 140,000. Data analysis was performed with Biopharma Finder 3.2 and intact protein spectra was automatically deconvoluted with Default-Auto Xtract.

### Liposome Preparation

Lipids in chloroform or in chloroform/methanol (2:1) in the case of mixtures containing PI(4)P were mixed at the desired molar ratio and the solvent was removed in a rotary evaporator. Lipid films were hydrated in 50 mM HEPES pH 7.4, 120 mM K acetate (HK buffer, which was degassed before use) to reach a concentration of 2 mM. The suspensions were then frozen in liquid nitrogen and thawed in a water bath (37°C) five times. Liposomes were extruded through 0.1 µm pore size polycarbonate filters using hand extruder (Avanti Polar Lipids) and were used within 1-2 days.

### Full length VAP-A reconstitution into proteoliposomes

VAP-A^FL^ WT or sL1/2 were reconstituted by detergent removal using BioBeads as described previously in lipid mixture containing egg PC and brain PS (95/5 mol%) for cryo-EM experiment or eggPC, brainPS, DGS NTA(Ni), cholesterol (93/5/2/10 mol%) for *in vitro* lipid transfer assay. As previously described in ref. (Mora et al., 2021), DDM solubilized VAP-A was mixed at room temperature with liposomes solubilized in Triton X100 (detergent/lipid ratio of 2.5 w:w), in 50 μL during 15 min. Thereafter, detergent was removed by successive additions of wet BioBeads, at BioBeads/detergent ratio of 10 w:w (for 2 h), 20 w:w (for 1 h), 20 w:w (for 1 h) at room temperature. All these steps were done in a 2-mL Eppendorf tube under gentle mixing with a small magnetic stir bar. The reconstitution buffer contained 1 mg/mL lipid, 2.5 mg/mL Triton X100 in HK. Protein was added at LPR ranging from 70 to 2800 mol/mol depending on experiment. All the reconstitutions were tested by flotation assay. Reconstitutions of VAP-A sL1/2 for cryo-EM experiment were performed with addition of 400 mM KCl to prevent the formation of multilayered vesicles and increase the homogeneous distribution of proteins in vesicles. After reconstitution, proteoliposomes were kept at 4°C and used within 2 days.

### Cryo-EM experiments

Lacey carbon 300 mesh grids (Ted Pella, USA) were used in all cryo-EM experiments. Blotting was carried out on the opposite side from the liquid drop and plunge frozen in liquid ethane (EMGP, Leica, Germany). Cryo-EM images were acquired with a Tecnai G2 (Thermofisher) Lab6 microscope operated at 200 kV and equipped with a 4k×4k CMOS camera (F416, TVIPS). Image acquisition was performed under low dose conditions of 10 e^-^/Å^2^ at a magnification of 50,000× with a pixel size of 2.1 Å/px.

### Formation of in vitro MCS

VAP-A^FL^/N-PH-FFAT MCS were formed directly on the cryo-EM grid at room temperature according to (Mora et al., 2021). VAP-A proteoliposomes were diluted at 0.05 mg/mL in HK buffer. Lipid nanotubes containing C24:1 Galactosyl(β) Ceramide, egg PC, brain PS, brain PI(4)P (80/10/ 5/5 mol%) were diluted at 0.15 mg/mL and mixed with N-PH-FFAT at LPR 70 mol/mol. A 2 μL drop of N-PH-FFAT tubes was immediately loaded onto the grid and left to incubate for ∼1 min, followed by the addition of 2 μL VAP-A^FL^ WT of sL1/2 proteoliposomes. After 30 sec incubation, the grid was blotted and immediately flash frozen.

### Liposome flotation assay

Liposomes containing egg PC, brain PS, liver PE, Rhodamine-PE and PI(4)P (62/5/17/2/4%) (1 mM) and N-PH-FFAT (1 µM) were incubated for 5 min in HKM buffer at room temperature in a Beckman polycarbonate centrifuge tube. 1 µM of VAP-A^His^ was added with or without preincubation with FFAT peptide 5 min at room temperature (Proteogenix Oberhausbergen, France WCSGKGDMSDEDDENEFFDAPEIITMPENLGH). The 150 µL mixture was incubated 10 min at 37°C under agitation and mixed with HK buffer supplemented with 60% sucrose (100 µL) and then overlaid with an intermediate sucrose solution made of HK buffer with 24% sucrose (200 µL) and a layer (50 µL) of HKM buffer. After centrifugation at 55,000 rpm in a TLS 55 (Beckman) swing rotor for 90 min at 25°C, three fractions (bottom, medium, and top) of 250, 150 and 100 µl were collected and analyzed by SDS-PAGE using Sypro-Orange staining.

### In vitro lipid transfer assay

PI(4)P-transfer assays were performed as described previously using purified recombinant proteins and liposomes mimicking ER and Golgi membranes (Mesmin et al., 2013). All measurements were carried out in a Jasco FP-8300 spectrofluorimeter using a cylindrical quartz cuvette (600 mL) equilibrated at 37°C and equipped with a magnetic bar for continuous stirring. Golgi-like donor (Ld) liposomes were prepared with egg PC, liver PE, liver PI, brain PS, brain PI(4)P (64/17/8/5/4 mol%), supplemented with 2 mol % Rho-PE and filter-extruded at 0.1 µm pore-size. ER-like acceptor VAP-A^FL^ proteoliposomes (La) were reconstituted at LPR 1300 mol:mol in egg PC, brain PS, DGS NTA(Ni) (93/5/2 mol%), supplemented with 10 mol% cholesterol. The sample initially contained NBD-PH-FAPP1 (100 nM) and Ld (250 µM) in HKM buffer. La (250 µM) containing VAP-A^FL^ (150 nM), OSBP (0.1 µM) and Sac1^His^ (0.1 µM) were then sequentially added at the indicated times. PI(4)P transport was followed by measuring the NBD emission measured at 530 (±10) nm upon excitation at 460 (±1) nm.

### CRISPR/Cas9 Knock-out

Ribonucleoprotein complexes were prepared by mixing 1.2 µL sgRNA (1 µM) and 1.7 µL of recombinant HiFi Cas9 Nuclease (62 µM) (IDT DNA) in phosphate-buffered saline (PBS) (5 µL total volume). Upon incubation (20 min at room temperature), the complexes were then mixed with 2 µL of Cas9 Electroporation Enhancer (IDT DNA) and 10^6^ RPE1 cells before electroporation using the Amaxa Nucleofector device. After 48 h, single cell seeding was performed in 96 well-plates. Edition of each clone was analyzed by sequencing and Western blot using Rabbit anti-VAP-A, VAP-B, VPS13A, PTPIP1 (RMDN3) and α-Tubulin primary antibody (Atlas antibodies) and HRP-conjugated secondary antibodies (Jackson ImmunoResearch Labs). 5’-3’ sequences of crRNA used in this study (sgRNA : crRNA and trRNA complex. TrRNA was determined by IDT DNA to adapt to Cas9 Nuclease): VAP-A, TGAAGACTACAGCACCTCGCCGG; VAP-B, TGAAGACTACAGCACCACGTAGG; VPS13A, GACGTGTTGAACCGGTTCTTGGG; and PTPIP51, TGAAGTCTGC GTATAGTCCAGGG.

### Cell staining and imaging

For immunofluorescence, cells cultured in on μ-slides (Ibidi) were washed once with PBS then fixed with paraformaldehyde (3%) and glutaraldehyde (0.1%) in PBS for 10 min. Fixed cells were soaked in PBS supplemented with NH4Cl (50 mM) for 10 min, and then permeabilized in permeabilization buffer (Saponin 0.05 %, BSA 0.2 %, PBS) during 45 min at RT°. Cells were then immuno-labeled with primary antibodies (rabbit anti-OSBP or CERT (Atlas antibodies), mouse anti-TOM20 (Santacruz) or sheep anti-TGN46 (Biorad), diluted in permeabilization buffer) for 2 h at RT°, washed 3 times with permeabilization buffer, further incubated with secondary Alexa Fluor-conjugated antibodies (Invitrogen; diluted in permeabilization buffer) for 30 min at RT°, and rinsed. For plasma membrane staining, Wheat Germ Agglutinin (WGA) coupled to Alexa Fluor 647 (Invitrogen) in PBS was apply at 5 µg/mL on fixed unpermeabilized cells for 10 min before 2 washes in PBS. Mitotracker Red (Invitrogen) staining was performed at 500 nM on living cells in growth medium during 30 min at 37°C, and then washed twice.

Confocal images were acquired using a Leica SP8 STED 3X (Leica Microsystems, Nanterre) equipped with a pulsed white light laser. All images were acquired through a 63x /1.4 NA Oil objective using the LAS X software (Leica Microsystems). Confocal images have a ∼100 nm pixel size.

### Mitochondria purification

Mitochondria with MAM were isolated according to (Wieckowski et al., 2009). 4 million of RPE1 cells KO for VAP-A stably expressing EGFP-VAP-A WT or sL1/2 were collected and pelleted at 600×g for 5 min in PBS. Cells were then resuspended in 1 mL of PBS supplemented with CaCl2 1.2 mM and MgCl2 1.06 mM, and centrifugated at 600×g 5 min at 4°C. Cell lysis was then performed in 2 mL of ice-cold Buffer-1 (225 mM Mannitol, 75 mM succrose, 0.1 mM EDTA and 30 mM Tris HCl pH 7.4) using a pre-chilled 2-mL Dounce homogenizer. Homogenization steps were carried out at 4°C. Cell disruption was controlled under a microscope: after ∼500 passes of the pestle, >80% of cells were broken. At this step, total GFP-fluorescence per tube was measured using a Fusion FX imager (Vilber). The lysate was then centrifuged twice at 600×g for 5 min at 4°C in order to eliminate unbroken cells and nuclei. The supernatant was centrifuged at 7000×g for 10 min at 4°C, and the resulting the pellet was resuspended in 1 mL of ice-cold Buffer-2 (225 mM Mannitol, 75 mM sucrose and 30 mM Tris HCl pH 7.4) before a new centrifugation step at 7000×g 10 min, 4°C. The pellet containing mitochondria and MAM was resuspended in 2 mL of ice-cold Buffer-2 and GFP-fluorescence was measured again. This suspension was finally centrifuged for 10 min at 10,000×g 4°C and crude mitochondrial pellet was resuspended in RIPA buffer for Western blot analysis to control mitochondria enrichment.

### Mitophagy assay

EGFP-Microtubule-associated protein light chain 3 (LC3) was used to monitor autophagy by visualizing its recruitment to autophagosomes. RPE1 WT or KO for VAP-A were transfected with EGFP-LC3 and plated in µ-slide (Ibidi) for 24 h. Cells were then treated with DMSO or Carbonyl Cyranide m-Chlorophenylhydrazone (CCCP) and proteases inhibitors (E64D and Pepstatin A 10 µM) for 2h in growth medium. The Last 30 min of treatment included Mitotracker Red labelling. After two PBS washes, cells were fixed with 4% PFA and 0.1% glutaraldehyde before imaging.

### CTL synthesis

Cholesta-5,7,9(11)-trien-3β-ol (cholestatrienol, CTL) was synthesized in three steps from 7-dehydrocholesterol. For 7-dehydrocholesteryl-3-o-β-acetate synthesis, 7-dehydrocholesterol (1.00 g, 2.60 mmol) was refluxed at 140 ℃ in 5 mL acetic anhydride (52 mmol) until complete dissolution (*i.e.*, approximately 1.5 h). After removal of acetic anhydride *in vacuo*, 10 mL distilled water and 10 mL heptane were added. The organic phase was washed two times with water then brine, dried over magnesium sulfate, filtered and concentrated *in vacuo*. The crude product was purified by column chromatography on silica gel using a gradient of heptane and ethyl acetate (from 100/1 to 70/30). Dehydrocholesteryl-3-o-β-acetate (660 mg, 60% yield) was isolated as white crystals. Next, for Cholesta-5,7,9(11)-trien-3β-yl acetate synthesis, a solution of dehydrocholesteryl-3-o-β-acetate (460 mg) in ethanol (27 mL), was heated to reflux, and then mercuric acetate (1 g), dissolved in a mixture of ethanol (16 mL) and acetic acid (550 mL) was added. The reaction mixture was refluxed for 30 min, then stirred at room temperature for 16 h. The solvents were evaporated to dryness, and after addition of chloroform (10 mL), the solution was filtered through Celite. The organic phase was washed successively with a saturated solution of sodium carbonate, water and brine, then dried over magnesium sulfate, filtered and concentrated *in vacuo*. The crude product was purified by column chromatography on silica gel using a gradient of heptane and ethyl acetate (from 100:1 to 70:30) to give pure cholesta-5,7,9(11)-trien-3β-yl acetate (230 mg, 50% yield) as white crystals. Finally, Cholesta-5,7,9(11)-trien-3β-ol was obtained following the procedure described by (Modzel et al., 2017) and was ultimately purified by column chromatography on silica gel using a gradient of heptane and ethyl acetate (from 100:1 to 50:50) then precipitated in acetonitrile to give a pure white solid (61 mg, 30% yield) that was stored in the dark under argon, at –80 °C.

### CTL imaging

Cells were seeded in µ-slides 35mm dishes (Ibidi) and transiently transfected with TagBFP-bGalT1 for 24 h. For CTL labelling, cells were incubated with CTL in complex with MCD (1:5 ratio) for 1 min at 37°C, rapidly washed three times with growth medium. For imaging, dishes were mounted in a stage chamber set at 37°C (Okolab) and cells were observed using an IX83 inverted microscope (Olympus) equipped with an iXon3 blueoptimized EMCCD camera (Andor) and an UPlanSApo 60X/1.35 Oil objective (Olympus). CTL was imaged using Semrock BrightLine filters (320/40 nm band-pass filter, 347 nm dichroic beam splitter, and 390/40 nm band-pass filter).

### Widefield time-lapse imaging

Cells were seeded in µ-slides 35mm dishes (Ibidi) and placed in an appropriate initial volume of growth medium so that homogenous mixing of SWG (50 nM) (freshly diluted before use) was achieved upon subsequent additions during the time course measurements. Widefield time-lapse microscopy was performed using an Olympus IX83 inverted microscope equipped with a Z-drift compensator, a scanning stage SCAN IM (Marzhauser), and an iXon3 camera (Andor). Cells plated were mounted in a stage chamber set at 37°C (Okolab). TagBFP, EGFP, and mCherry were observed using Chroma fluorescence filter sets (ref. 49000, 39002, 39010). Multidimensional acquisition and analysis were performed with MetaMorph software (Molecular Devices).

### PEG fusion assay

50,000 cells expressing mitochondrially targeted CFP were cultured overnight on µ-slides 35mm (Ibidi) dishes with 50,000 cells expressing mitochondrially targeted mCherry. The next morning, cells were fused for 60s in 1 mL of 50% PEG 1500 (Roche). Cells were washed 4 times and incubated for 16 h in growth medium containing 20 µg/mL cycloheximide before fixation. Mitochondria fusion was analyzed by confocal microscopy and quantified using Pearson’s correlation coefficient measurement.

### Fluorescence Recovery After Photobleaching (FRAP) assay

For FRAP assay, cells were maintained at 37°C in growth medium. FRAP were carried out using a Nikon Eclipse Ti inverted microscope equipped with an UltraVIEW VoX spinning disc imaging system (PerkinElmer) driven by Volocity software. Images were acquired using a 100× oil-immersion objective (Nikon CFI Plan Apochromat 100×/1.4). Photobleaching was performed on circular areas of ∼4 μm^2^ within cell perinuclear regions. Fluorescence in these areas was then recorded every 0.5 s and normalized to the initial intensity.

### Stimulated Emission Depletion microscopy (STED)

Cells were transfected with pSNAP-tag or pHTN HaloTag® plasmids using the Amaxa nucleofector device (Lonza), and plated on glass coverslips. Cells were labelled 24 h after transfection with 3 µM SNAP-Cell TMR STAR (New England BioLabs) or 0.2 µM Jenelia Fluor 646 HaloTag®-Ligand (Promega) in growth medium for 30 min at 37°C. Cells were then washed twice to remove excess dye and further incubated for 30 min at 37°C in growth medium. Finally, cells were washed in PBS, fixed with PFA 4%/glutaraldehyde 0.1%, washed again in PBS and quicky in water before mounting in Abberior Mount Solid Antifade (Abberior GmbH). STED images were acquired using a Leica SP8 STED 3X (Leica Microsystems, Nanterre) equipped with a pulsed white light laser as an excitation source and the 775 nm pulsed laser as depletion light source. All images were acquired through a 100x/1.4 Oil objective using the LAS X software (Leica Microsystems). The TMR Star was imaged with a 561 nm excitation wavelength and the Jenelia Fluor 646 was imaged with a 633 nm excitation wavelength. STED images have a 15 nm pixel size. All images were deconvolved with Huygens Core version 21.04/Huygens Remote Manager version 3.7 (Scientific Volume Imaging), using the CMLE algorithm.

### Transmission Electron Microscopy (TEM)

For TEM, cells were fixed with 1.6% glutaraldehyde in 0.1 M phosphate buffer, rinsed in 0.1 M cacodylate buffer, and postfixed for 1 h in 1% osmium tetroxide and 1% potassium ferrocyanide in 0.1 M cacodylate buffer to enhance membrane staining. The cells were then rinsed in distilled water, dehydrated in alcohols and lastly embedded in epoxy resin. Contrasted ultrathin sections (70 nm) were analyzed under a JEOL 1400 transmission electron microscope mounted with a Morada Olympus CCD camera.

### Mass Spectrometry

A modified Bligh and Dyer was used for lipid extraction. One million cells were collected and pelleted in an Eppendorf tube and resuspended in 200 µL of pre-cooled water, 500 µL of methanol and 250 µL of chloroform. The mixture was vortexed for 10 min before a new addition of 250 µL of water and 250 µL of chloroform. The mixture was vortexed again for 10 min and finally centrifuged (3000 rpm, 15 min at 4°C). Then, 400 µL of the organic phase was collected in a new glass tube and dried under a stream of nitrogen. The dried extract was resuspended in 60 µL of methanol/chloroform (1:1 v/v) and transferred in an injection vial.

Reverse phase liquid chromatography was selected for separation with an UPLC system (Ultimate 3000, ThermoFisher). Lipid extracts were separated on an Accucore C18 (150×2.1, 2.5 µm) column (ThermoFisher) operated at 400 µL/min flow rate. The injection volume was 3 µL. Eluent solutions were ACN/H_2_O (1:1 v/v) containing 10 mM ammonium formate and 0.1% formic acid (solvent A) and IPA/ACN/H_2_O (88:10:2 v/v) containing 2 mM ammonium formate and 0.02% formic acid (solvent B). The step gradient of elution was in %B: 0 min, 35%; 0‒4 min, 35 to 60%; 4‒8 min, 60 to 70%; 8‒16 min, 70 to 85%; 16‒25 min, 85 to 97%; 25‒25.1 min 97 to 100% B, 25.1‒31 min 100% B and finally the column was reconditioned at 35% B for 4 min. The UPLC system was coupled with a Q-exactive Mass Spectrometer (Thermofisher); equipped with a heated electrospray ionization (HESI) probe. This spectrometer was controlled by Xcalibur software (version 4.1.31.9.) and operated in electrospray positive mode.

For cardiolipin analyzes lipid extracts were separated on a Cortex C8 (150×2.1, 2.7 µm) column (Waters) operated at 400 µL/min flow rate (column temperature 45°C). The injection volume was 3 µL of diluted lipid extract. Eluent solutions were ACN/H_2_O (6:4 v/v) containing 10 mM ammonium formate and 0.1% formic acid (solvent A) and IPA/ACN/H_2_O (88:10:2 v/v) containing 2 mM ammonium formate and 0.02 % formic acid (solvent B). The used step gradient of elution was: 0‒5 min 60% B, 5‒12.5 min 60 to 74% B, 12.5‒13 min 70 to 99% B, 13‒17 min 99% B, 17‒17.5 min 99 to 60% B and finally the column was reconditioned at 60% B for 2.5 min. In this case the Mass Spectrometer was operated in electrospray negative mode.

Data were acquired with dd-MS2 mode. MS spectra were acquired at a resolution of 70,000 (200 m/z) in a mass range of 250−1200 m/z or 1000−1600 m/z for CL analyzes with an AGC target value of 1e6 and a maximum injection time of 250 ms. 15 most intense precursor ions were selected and isolated with a window of 1 or 1,5 m/z and fragmented by HCD (Higher energy C-Trap Dissociation) with normalized collision energy (NCE) of 25 and 30 eV or 20, 30 and 40 eV for CL analyses. MS/MS spectra were acquired in the ion trap with an AGC target 1e5 value, the resolution was set at 35 000 at 200 m/z combined with an injection time of 80 ms.

Data were reprocessed using Lipid Search 4.2.21 (ThermoFisher). The product search mode was used and the identification was based on the accurate mass of precursor ions and MS2 spectral pattern. The parameters for the ion identification and alignment were set up as follow: ion precursor tolerance 5 ppm; ion product tolerance 5 ppm; ion searched [H+], [NH4+] and [Na+] or [H-] for CL analyzes; alignment retention time tolerance 0.4 min; ID quality filters A, B and C.

### MD simulations of dimer models

The dimer models of VAP-A WT and sL1/2 were built with AlfaFold2 tool (Jumper et al., 2021). The VAP-A transmembrane helix was removed. MD simulations were performed with GROMACS 2021.3 (https://www.gromacs.org/) with the CHARMM36 force field. The dimer was centered in the cubic box, at a minimum distance of 2 nm from box edges. Solvent molecules were added in the coordinate file. The TIP3P water model configuration was used. Na and Cl ions were added to neutralize the simulation box, at a minimum concentration of 120 mM. The total number of atoms (protein + solvent + counterions) was 462,599 for the VAP-A WT dimer model and 258,855 for sL1/2 dimer model.

Energy minimization was performed using the steepest descent minimization algorithm for the subsequent 50,000 steps. A step size of 0.01 nm was used during energy minimization. A cutoff distance of 1 nm was used for generating the neighbor list and this list was updated at every step. Long-range electrostatic interactions were calculated using the particle mesh Ewald summation methods. Periodic boundary conditions were used. A short 100 ps NVT equilibration was performed. During equilibration, the protein molecule was restrained. All bonds were constrained by the LINear Constraints Solver (LINCS) constraint algorithm. A short 100 ps NPT equilibration was performed similar to the NVT equilibration. During the production run, the V-rescale thermostat and Parrinello–Rahman barostat stabilized the temperature at 300 K and pressure at 1 bar, respectively. The simulations were performed for 1 µs for each system and coordinates were saved every 100 ps. The MD analyzes of the Root Mean Square Fluctuation (RMSF) were performed using GROMACS utilities. Trajectories have been centered and fitted on the coiled-coiled domain of VAP-A. Movies and images have been generated with VMD and PyMOL 2.0 software, respectively.

### QUANTIFICATION AND STATISTICAL ANALYSIS

In cryo-EM images, the lipid bilayer of the vesicles or of the ribbons is identifiable and serves as landmarks for distance measurements. In the plot Fig. 1g the lengths of the extramembraneous domains of VAP-A are reported from the electron densities extending at the largest distances from the membrane.

For EGFP-VAP-A enrichment at the TGN area, a TGN mask was applied on the GFP channel and a ratio between the average fluorescence in this area and the average fluorescence in the rest of the cell (*i.e.*, cell minus TGN area) is determined. For TGN/cytosol ratios of OSBP, P4M, CTL and CERT, two regions of interest (ROIs) of the same area were drawn in the TGN and in the cytosol. The average fluorescence was determined for each ROI and the ratio was then calculated. In all superplots illustrating image quantification, each point represents one cell analyzed, and each experiment was performed 3 times independently. Statistical analyzes were done in Prism * p < 0.1, **p < 0.01, *** p < 0.001, **** p < 0.0001. The scale bars in widefield, confocal and STED images are 10 µm, and 2 µm in zoomed images. The images were shared, treated and edited using OMERO.web, OMERO.iviewer, OMERO.figure from an OMERO image database from the Université Côte d’Azur and EMBRC-France OMERO Database.

Colocalization quantifications were done using Coloc 2 Pearson test on Fiji software (ImageJ, NIH). Kymographs were generated using the Metamorph software (Molecular Devices) from a rectangle drawn on the image stack and projected across time of the complete time series. Scan line quantifications were generated using Fiji software. For mCherry-D4H quantification at the plasma membrane a mask of WGA was applied on the D4H channel, normalized by the total intensity in the field and by the control condition; one point represents one field analyzed; experiments were repeated 3 times independently with 8 fields analyzed per experiment.

To quantify mitochondria morphology, maximum intensity projection images from confocal z-series were processed automatically through a custom pipeline in CellProfiler (Broad Institute). Images were first background corrected and a gaussian filter was applied using the *CorrectIllumination* module. Mitochondria were then accentuated and binarized using the *EnhanceOrSuppressFeatures* and *Threshold* modules. Thereafter, images were cleaned using the *Morph* module, and mitochondria (objects) were identified using the *IdentifyPrimaryObject* module. Finally, the *MeasureObjectSizeShape* module enabled measuring objects shape, notably according to the form factor [(4×π×Area)/Perimeter²]. Values were then ranked in a histogram where the number of mitochondria for each form factor value was normalized to the total mitochondria count. Data was further normalized to the mean maximum count of the WT condition. For TEM images quantification, mitochondria width was measured in Fiji (Image J, NIH) by drawing straight lines. One point represents one mitochondrion analyzed.

## SUPPLEMENTAL VIDEO TITLE AND LEGEND

**Movie S1. Mitochondria dynamics in RPE1 WT and VAP-A KO cells. Related to Figure S4b.** Time-lapse confocal imaging of Mitotracker-labelled cells showed in Figure S4 featuring mitochondrial fusion and fission events. In the VAP-A KO condition, the red arrow shows two spherical mitochondria close to each other but not fusing.

