## Supplemental information for "VAP-A intrinsically disordered regions enable versatile tethering at membrane contact sites"

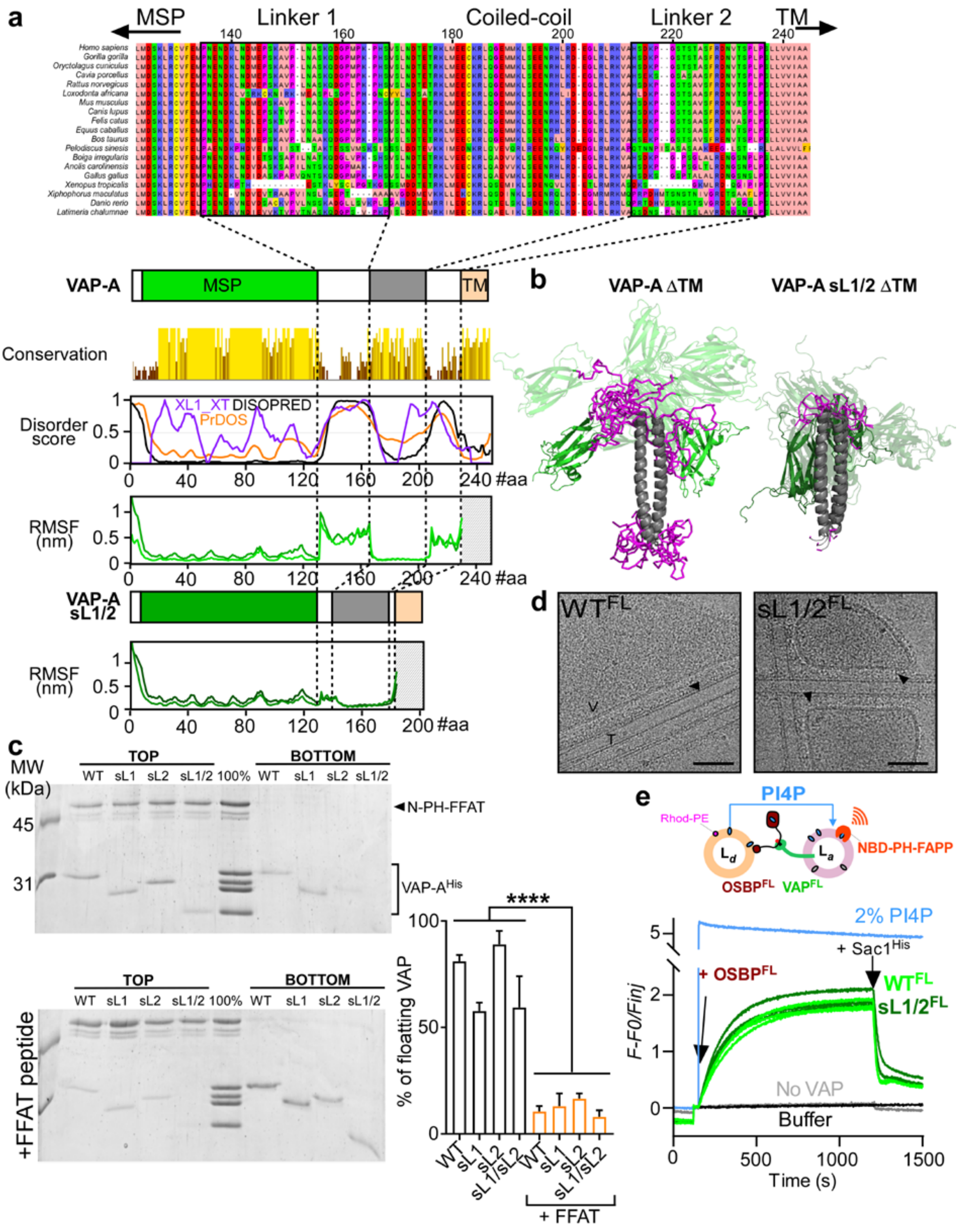


**Figure S1. Intrinsically disordered regions in VAP-A. Related to figure 1. a**, Clustal-O alignment of VAP-A sequences from vertebrates. The conservation score (yellow scale) is proportional to the bars' height. Intrinsically disorder prediction from different algorithms

(XL1_XT, DISOPRED and PrDOS) suggests that linkers 1 and 2 are unstructured. The RMSF is calculated from molecular dynamics simulations with VAP-A ΔTM. The two lines represent RMSF of each monomer of the dimeric protein. Linker regions show greater fluctuation sizes. By contrast, VAP-A short linkers (sL1/2) show fluctuations indistinguishable from the rest of the structure, as expected. **b**, Snap-shots from molecular dynamics simulations of VAP-A WT or sL1/2 (ΔTM), built with the AlfaFold2 tool, showing that the movements of the MSP are more restricted with the mutant construct. **c**, Floatation assay. N-PH-FFAT bound to PI(4)P liposomes was incubated with VAP-$A^{His}$ WT or sL mutant constructs. The suspension was adjusted to 30% w/v sucrose and overlaid with two cushions of decreasing sucrose density. After ultracentrifugation, the top and bottom fractions were collected and analyzed by SDS-PAGE and Sypro-orange staining. Proteins bound to liposomes were retrieved on the top fraction. Bottom; a peptide encompassing the FFAT motif of OSBP was added as a competitor. Right; quantification of the fraction of VAP-$A^{His}$ bound to N-PH-FFAT. Data are mean of 3 independent experiments ± SEM. **d**, Representative cryo-EM images of *in vitro* reconstituted MCS between proteoliposomes containing VAP-A WT or sL1/2 (LPR 70 mol/mol) and N-PH-FFAT bound to galactosylcerebroside nanotubes dopped with PI(4)P. Protein densities at MCS are indicated by arrowheads. V: vesicle; T: tube. Bar: 50 nm. **e**, Real-time measurement of OSBP-mediated PI(4)P transfer *in vitro*. NBD-PH-FAPP1 (250 nM) was mixed with donor liposomes (Ld; 250 µM lipids containing 4 mol% PI(4)P and 2 mol% Rhod-PE) and acceptor liposomes (La; 250 µM lipids containing 2 mol% DGS-NTA(Ni) and 150 nM VAP-A reconstituted at LPR 1300 mol/mol). At t=150s, OSBP (100 nM) was added to the mix. Thereafter, $Sac1^{His}$ (100 nM) was added at t=1200s in order to hydrolyze PI(4)P transferred to La. The blue curve represents the signal for full equilibration of PI(4)P between Ld and La.

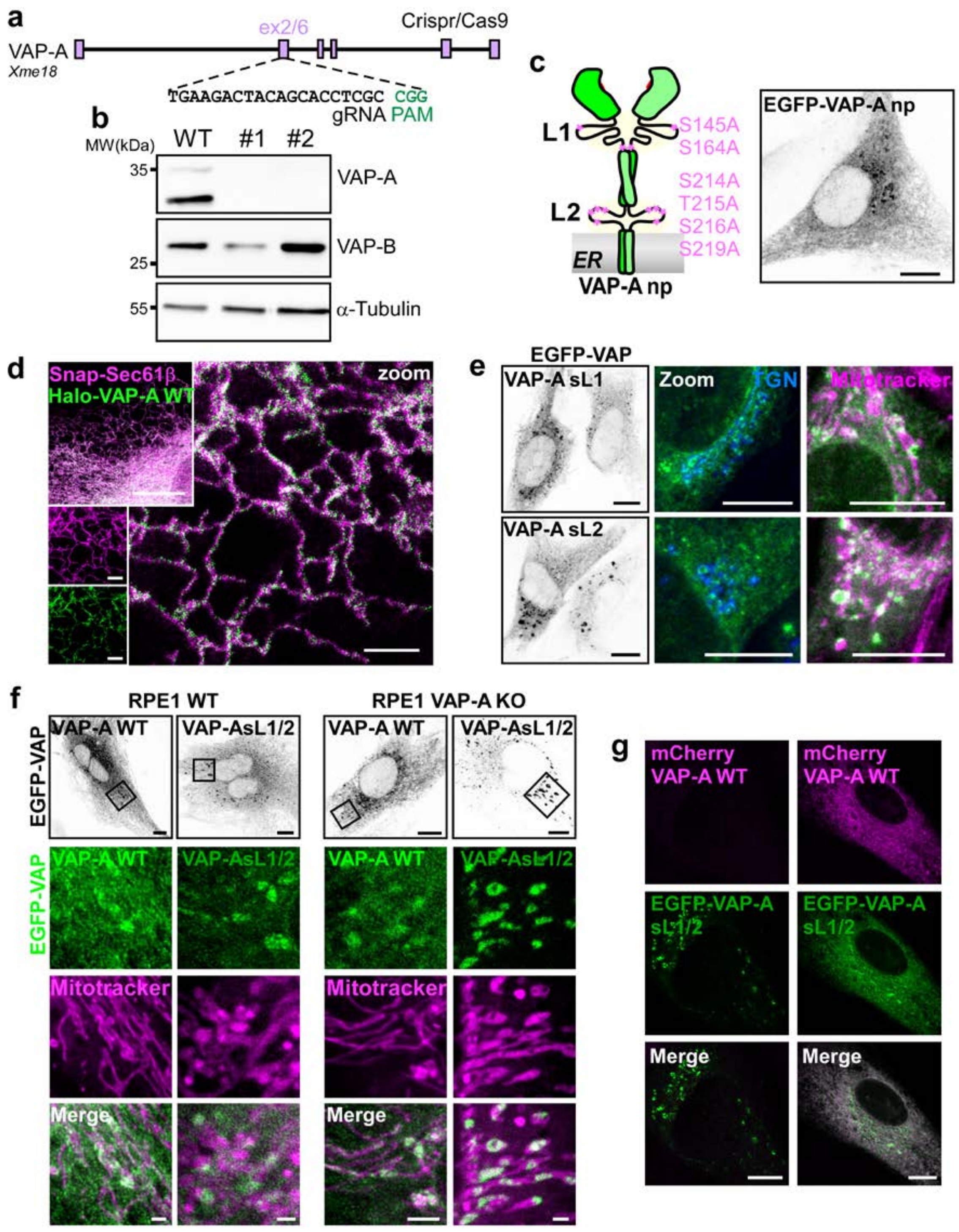


**Figure S2. CRISPR/Cas9-mediated KO and subcellular localization of VAP-A mutants. Related to figure 2. a**,**b**, CRISPR/Cas9 generation of RPE-1 VAP-A KO cells, targeting strategy and Western blot analyzes. **c**, Representative image of RPE-1 VAP-A KO cell expressing EGFP-VAP-A np, in which all phosphorylable residues present in the linkers were replaced by alanines. **d**, STED image showing colocalization between Halo-VAP-A WT and the ER labelled by Snap-Sec61β. **e**, Confocal images showing partial redistribution of EGFP-VAP-A sL1 or sL2 to ER–Mito MCS in RPE1 cells KO for endogenous VAP-A. **f**, Comparison of EGFP-VAP-A WT or sL1/2 expressed in WT RPE-1 or in VAP-A KO RPE-1. The relocation of VAP-A sL1/2 to mitochondria is attenuated in RPE1 containing endogenous VAP-A WT. **g**, The co-expression of mCherry-VAP-A WT with EGFP-VAP-A sL1/2 in VAP-A KO RPE1 cells redistributes EGFP-VAP-A sL1/2 away from ER–Mito MCS. Data from **f** and **g** suggest that the mutant is able to form heterodimer with WT forms.

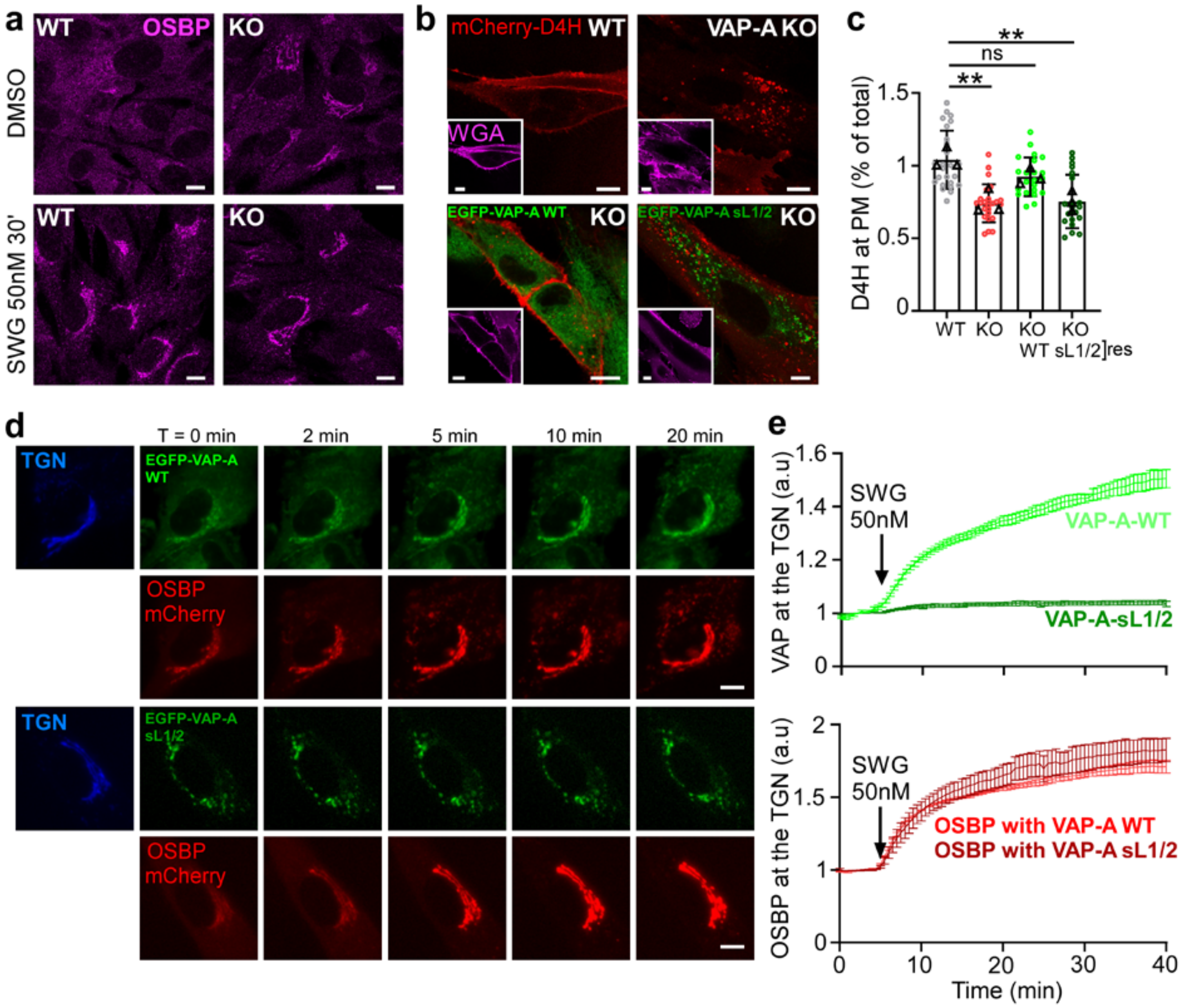


**Figure S3. Effects of OSBP inhibitor SWG treatment and cholesterol probe distribution. Related to figure 3. a**, Confocal images showing endogenous OSBP distribution in WT or VAP-A KO RPE-1 cells upon SWG (50 nM) treatment during 30 min. **b**,**c**, Cells were transfected with the cholesterol probe mCherry-D4H and labeled with the plasma membrane marker WGA-Alexa Fluor 647. The plot shows the amount of mCherry-D4H at the plasma membrane. Mean of 3 independent experiments ± SEM (24 fields were analyzed). **d**,**e**, Time-lapse imaging of VAP-A KO cells co-expressing EGFP-VAP-A WT or sL1/2, OSBP-mCherry, and the TGN marker TagBFP-βGalT1. OSBP and VAP-A WT, but not VAP-A sL1/2, are rapidly recruited to ER–Golgi MCS upon SWG (50 nM) addition. Graphs show the normalized mean fluorescence intensity of OSBP-mCherry or EGFP-VAP-A from TGN regions over the time course. Data are mean of 3 independent experiments ± SEM (n= 45 cells analyzed).

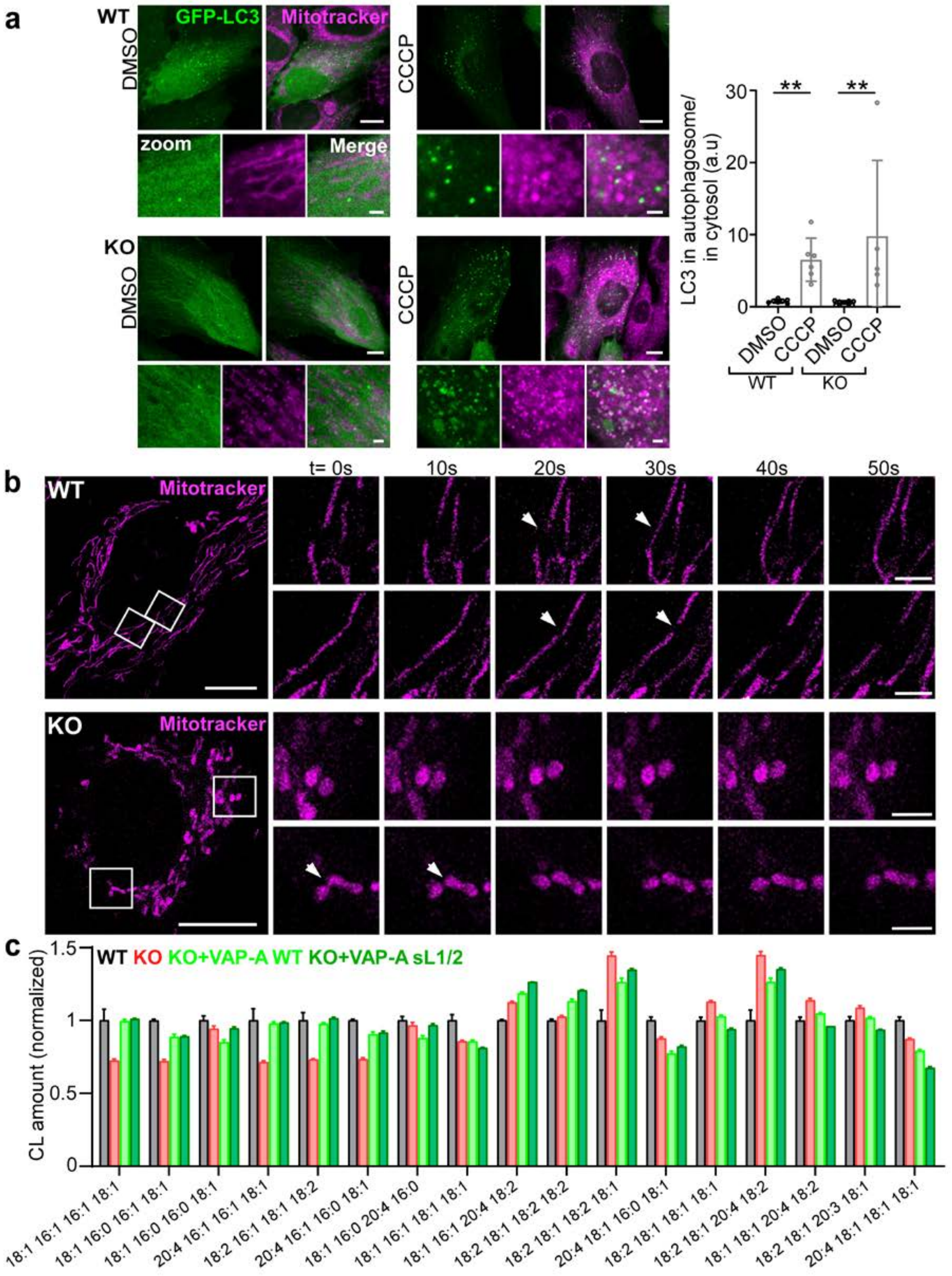


**Figure S4. VAP-A KO impairs mitochondrial fusion and CL metabolisms. Related to figure 4.**

**a**, Confocal images of RPE1 cells WT or KO for VAP-A expressing the autophagy marker GFP-LC3 in DMSO (control) or CCCP (mitophagy inducer) treatment conditions. The graph on the right is the quantification of the LC3 intensity at autophagosomes. **b**, Time-lapse confocal imaging of

Mitotracker featuring mitochondrial fusion and fission events (arrowheads). Fission, but no fusion, was observed in VAP-A KO cells. **c**, Quantification of CL species amount by LC-MS/MS. The relative areas are normalized to the control condition. Data are mean of 4 independent experiments ± SEM.

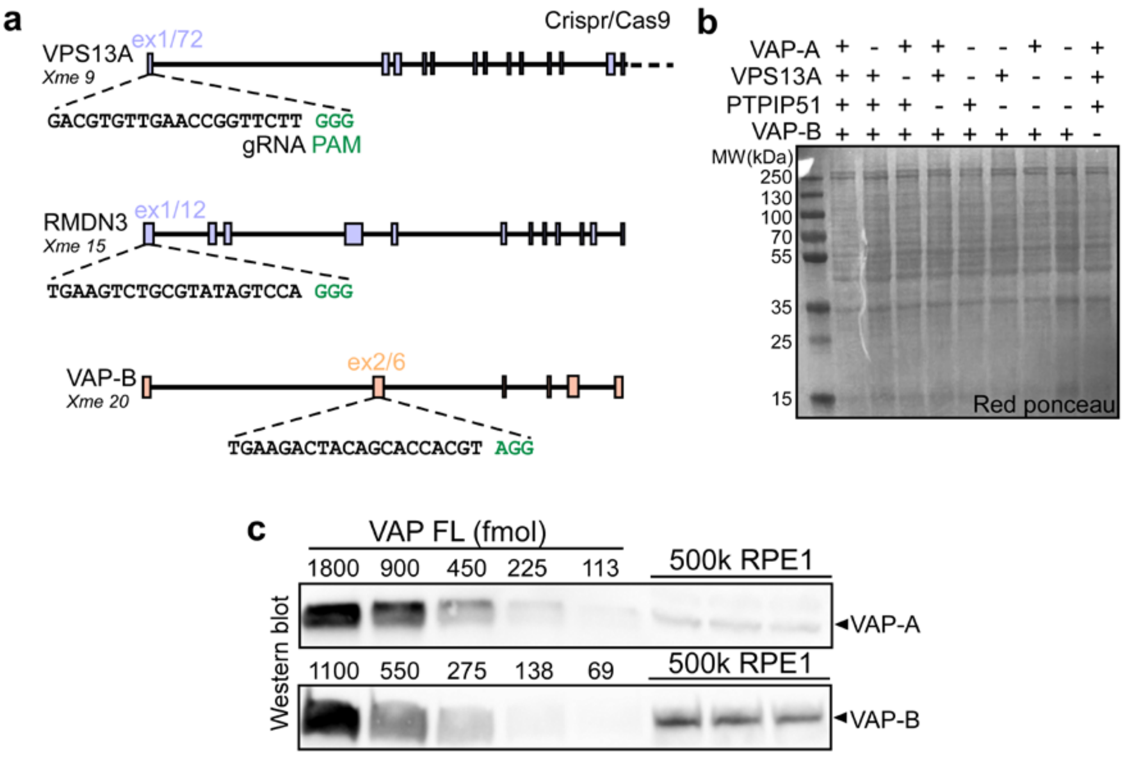


**Figure S5. Crispr/Cas9 KO strategy and endogenous VAP quantification in RPE1 cells. Related to figure 5. a**, Generation of RPE1 cell lines KO for VAP-A/B, PTPIP51 and/or VPS13A by CRISPR/Cas9. **b**, Red ponceau staining corresponding to the Western blot showed in fig. 5a. **c**, Quantification of endogenous VAP-A and VAP-B in RPE1 cells by Western blot. Standards are from known amounts of purified full-length VAP-A or VAP-B. Total lysate from 500.000 cells was loaded in each of the three last lanes. Proteins were then immunoblotted with anti-VAP-A (top) or anti-VAP-B (bottom). We measured that one RPE1 cell contains 122 fmol (~147,000 molecules) of VAP-A and 254 fmol (~306,000 molecules) of VAP-B.

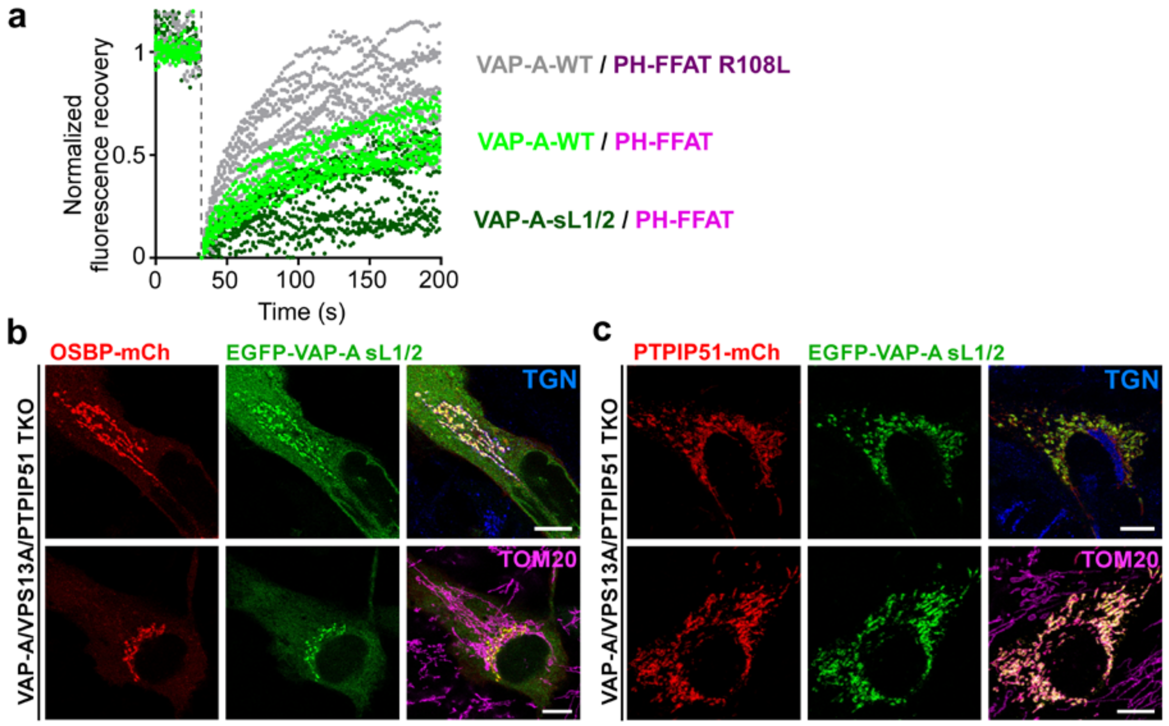


**Figure S6. FRAP experiment and subcellular distribution of EGFP-VAP-A sL1/2. Related to figure 6. a**, FRAP analysis performed on EGFP VAP-A WT or sL1/2 on RPE1 cells coexpressing PH-FFAT-mCherry WT or R108L, as indicated. Data are from 2 independents experiments (n≈10 cells analyzed). **b**,**c**, Transient expression of EGFP-VAP-A-sL1/2 with OSBP-mCherry or PTPIP51-mCherry in RPE-1 KO for VAP-A, VPS13A and PTPIP51.

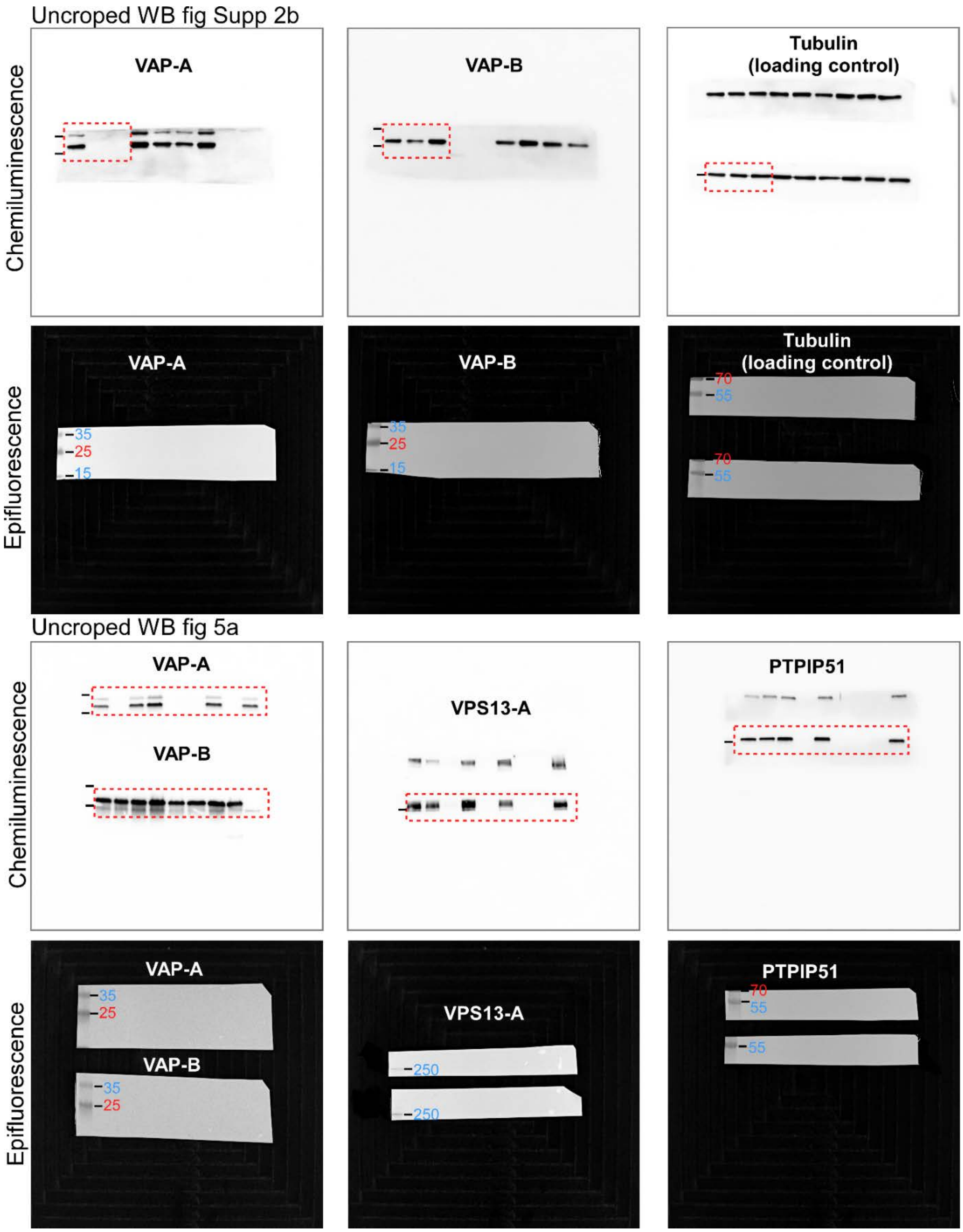


**Figure S7. Unprocessed gels.**
